# Cryo-EM resolves the structure of the archaeal dsDNA virus HFTV1 from head to tail

**DOI:** 10.1101/2024.12.09.627619

**Authors:** Daniel X. Zhang, Michail N. Isupov, Rebecca M. Davies, Sabine Schwarzer, Mathew McLaren, William S. Stuart, Vicki A.M. Gold, Hanna M. Oksanen, Tessa E.F. Quax, Bertram Daum

**Author notes:** Medical Sciences Doctoral Training Centre, Medical Sciences Division, University of Oxford, Oxford, UK. Biology Department, University of York, York, YO10 5DD, UK.

## Abstract

Outnumbering their hosts by at least a factor of 10, viruses are the most common biological entity on Earth, are major drivers of evolution, and greatly impact on the dynamics of our planet’s ecosystems. While viruses infecting bacteria and eukaryotes have been extensively studied, the viruses roaming the archaeal domain remain largely unexplored. In recent years, a growing number of archaeal viruses have been described, revealing a stunningly diverse range of morphologies that appear unique to archaea. Detailed structural studies are paramount to fully understand how archaeal viruses infect their hosts. However, no complete atomic models of archaeal viruses are available to date. Using electron cryo-microscopy, we investigated the structure of the archaeal virus Haloferax tailed virus 1 (HFTV1), which infects the halophile Haloferax gibbonsii LR2-5 originating from the Senegalese salt lake Retba. Through single particle analysis, we achieved near-atomic resolution for the entire set of HFTV1’s structural proteins, enabling the building of a full atomic model of the virion. Comparing the structures of DNA filled and empty capsids, we visualise structural changes occurring upon DNA ejection. By investigating the double-stranded DNA inside the capsid, we elucidate how the genome is spooled upon loading. Furthermore, our structure reveals putative cell-surface receptor-binding and catalytic roles of capsid turret, baseplate, and tail fibre proteins. Together, our data provide new insights into the mechanisms of HFTV1 assembly and infection, unveiling new perspectives on general rules of host-virus interactions in archaea and their evolutionary links to bacterial and eukaryotic viruses.

## Introduction

Viruses infecting microorganisms are abundant and diverse and those infecting archaea encompass an astonishing variety of morphotypes, such as spiral, bottle, and spindle-shaped viruses [1]. Of all different morphotypes, viruses with a helical tail and an icosahedral head containing a double-stranded DNA (dsDNA) genome are extremely successful. As such, they represent the predominant archetype of bacteria-infecting viruses and exist in abundance in almost all ecosystems of this planet. These tailed viruses (TVs) infect all major lineages of bacteria, drive cellular evolution, and have a major impact on ecosystems, such as the oceans, and are thus important for nutrient turnover [2–7].

Archaea are also infected by a diverse group of TVs (arTVs), which are evolutionarily related to the bacteria-infecting TVs (baTVs) and have been grouped together with them in the class *Caudoviricetes* [8,9], in which several new arTV-specific families have been established. Specifically, halophilic and methanogenic archaea belonging to the *Euryarchaea* are often infected by arTVs and encode arTV-like proviruses [9–18]. In addition, arTV-like proviruses are found in the genomes of other major classes of archaea, such as *Thaumarchaea*, *Aigarchaea*, and *Thermoplasmata* [19–22]. Comparison of arTV and baTV genomes suggests an ancient divergence, and arTVs represent an evolutionarily distinct group within the prokaryotic dsDNA virome [8]. The evolutionary link between the two groups is shown by their common virion architecture and conserved proteins, especially their major capsid proteins (MCPs) [23,24]. Based on the broad distribution of TVs, they are hypothesised to be a part of the virome of the last universal common ancestor (LUCA) [8,25].

dsDNA TVs are the most successful prokaryotic viruses known to us. The structure and infection mechanism of these viruses have been studied predominantly for bacterial virus members of the *Caudoviricetes*. Like baTVs, arTVs also have different tail types, which group them into podo-, myo- and siphovirus morphotypes [9].

Generally, the tails are involved in host recognition, binding, penetration of the cell envelope, and genome delivery into the host cell [26,27]. Some structural proteins of baTVs also have enzymatic activity, such as the peptidoglycan hydrolase activity of the tape measure proteins of mycobacteriophages [28]. However, the lack of full atomic structures for tape measure proteins precludes a full understanding of their function.

After genome delivery, the transcriptional program of the virus is activated, the genome replicated, and progeny particles are assembled in the cytoplasm. Most capsids of baTVs are icosahedral, but some are elongated (prolate) [29]. Usually, the capsid head is connected to the tail by a portal protein [29]. During virus maturation, this is the site where assembly of the head is initiated [29]. Heads consist of the MCPs, which require scaffolding proteins to obtain the correct capsid geometry [29]. Maturation of the empty capsid occurs by proteins that degrade the scaffolding protein.

The viral genome is packaged into the empty head in an energy-dependent manner by a terminase complex transiently docked to the portal vertex. Previous electron cryo-microscopy (cryo-EM) studies of baTVs have shown that dsDNA typically forms coaxial spools within viral capsids, in which the strands of DNA form ordered shells around an axis aligned with the portal [30–32]. These shells display increasing disorder towards the centre, possibly due to the inner layers being further from the capsid and therefore less influenced by the icosahedral structure, which may act as a guide for the more ordered outer layers.

While other types of genome spooling, such as concentric and toroidal organisations have been suggested, the coaxial spooling model is the most widely accepted model for dsDNA inside icosahedral capsids [32]. However, molecular dynamics studies, which can reproduce physical properties and packaging forces of the capsid, have suggested a more disordered organisation of DNA, often with multiple configurations [33]. These models have also revealed that the organisation of the DNA depends on the size and shape of the capsid; elongated icosahedrons have been simulated to display toroidal packaging in comparison to isometric icosahedrons, which typically display coaxial spooling [32].

After DNA packaging, the portal is plugged and connected to the tail. baTVs with short tails, such as podoviruses like phage T7, are assembled directly on the portal vertex, while the longer tails of myo- and siphoviruses are first assembled and connected later to the head [29].

The infection and assembly mechanisms of arTVs are less studied than those of baTVs. However, it was recently shown that the receptor for two arTVs is the S-layer protein [34,35], which is the main cell envelope component of archaea. We have shown that the siphovirus Haloferax tailed virus 1 (HFTV1) contacts its host *Haloferax gibbonsii* LR2-5 via its tail, but interestingly also with its head [15]. HFTV1 is hypothesised to bind the cell first reversibly by its head and then irreversibly via its tail, before ejecting its DNA into the host cell [35]. The structural details behind this unusual binding mechanism remain elusive.

Comparative genomics have indicated that arTVs possess MCPs with the same Hong Kong 97 (HK97) fold as is conserved amongst baTVs and eukaryotic herpesviruses [36]. Indeed, by electron cryo-tomography (cryo-ET) it was shown that the head of the arTV *Haloarcula sinaiiensis* tailed virus 1 (HSTV-1) consists of multiple MCPs with a HK97 fold, evolutionarily linking the arTVs with baTVs and herpesviruses (the realm *Dublodnaviria*) [23,24,37]. However, the complete structure of any arTV virion has not yet been resolved, and it is therefore unclear to what extent arTVs compare with baTVs of the class *Caudoviricetes*.

Here, we purified HFTV1 virions and studied their structure via cryo-EM. Through single particle analysis, we obtained near-atomic resolution maps of DNA-filled and empty virions, enabling us to assemble full atomic models of the virus pre and post DNA release - including all structural protein components. Comparing both structures enabled us to propose a model that describes the hitherto unknown host-cell adsorption and infection mechanism. Additionally, we were able to visualise the packaged dsDNA, revealing how the DNA is spooled inside the capsid.

Finally, our structure will aid in tracing the evolutionary history and structural diversity of the members of the class *Caudoviricetes* and to understand the common principles that underpin the global success of these widely distributed prokaryotic viruses.

## Results

### Structures of the HFTV1 virion pre and post DNA ejection

To determine the structure of the full HFTV1 virion, we purified virus particles, plunge-froze them on holey carbon grids and imaged them using a Titan Krios TEM equipped with either a K3 Summit or Falcon 4i direct electron detector. The resulting movies were corrected for beam-induced drift and CTF and high-resolution maps of the virus were reconstructed using RELION-4.0 and 5.0 [38,39].

The sample of purified virions contained DNA-filled particles, but also those that had already ejected their DNA genomes. The latter were easily distinguishable from their infectious counterparts in raw micrographs and during 2D classification (Supplementary Figure 1 & 2), allowing for the two classes of particles to be processed separately and thus the viral structure pre and post ejection to be compared.

The final map of the whole infectious DNA-filled virion had a global resolution of 4.39 Å (Cryo-EM Flowchart 1; Supplementary Figure 11). To obtain higher resolutions for its constituent parts, the head, turrets, portal, tail, baseplate and tail fibres were subjected to focused refinements employing symmetry operators pertaining to each sub-structure (Cryo-EM Flowchart 1; Supplementary Figures 11-13). This was a complex undertaking, due to symmetry mismatches that occurred across various protein-protein interfaces in the virus. Notwithstanding this, the combined data allowed for atomic model building of the entire virion, aided by ModelAngelo [40] (Supplementary Movie 1). HFTV1’s full atomic structure (Fig. 1a,b) revealed a virus with a length of 1,350 Å and an icosahedral head with a diameter of ∼600 Å. The head is decorated by up to 60 turrets, which project ∼90 Å above the surface of the capsid and thus increase its diameter to ∼780 Å. The tail has a length of 560 Å and terminates into a baseplate with three radiating tail fibres of ∼190 Å in length.

**Figure 1.**
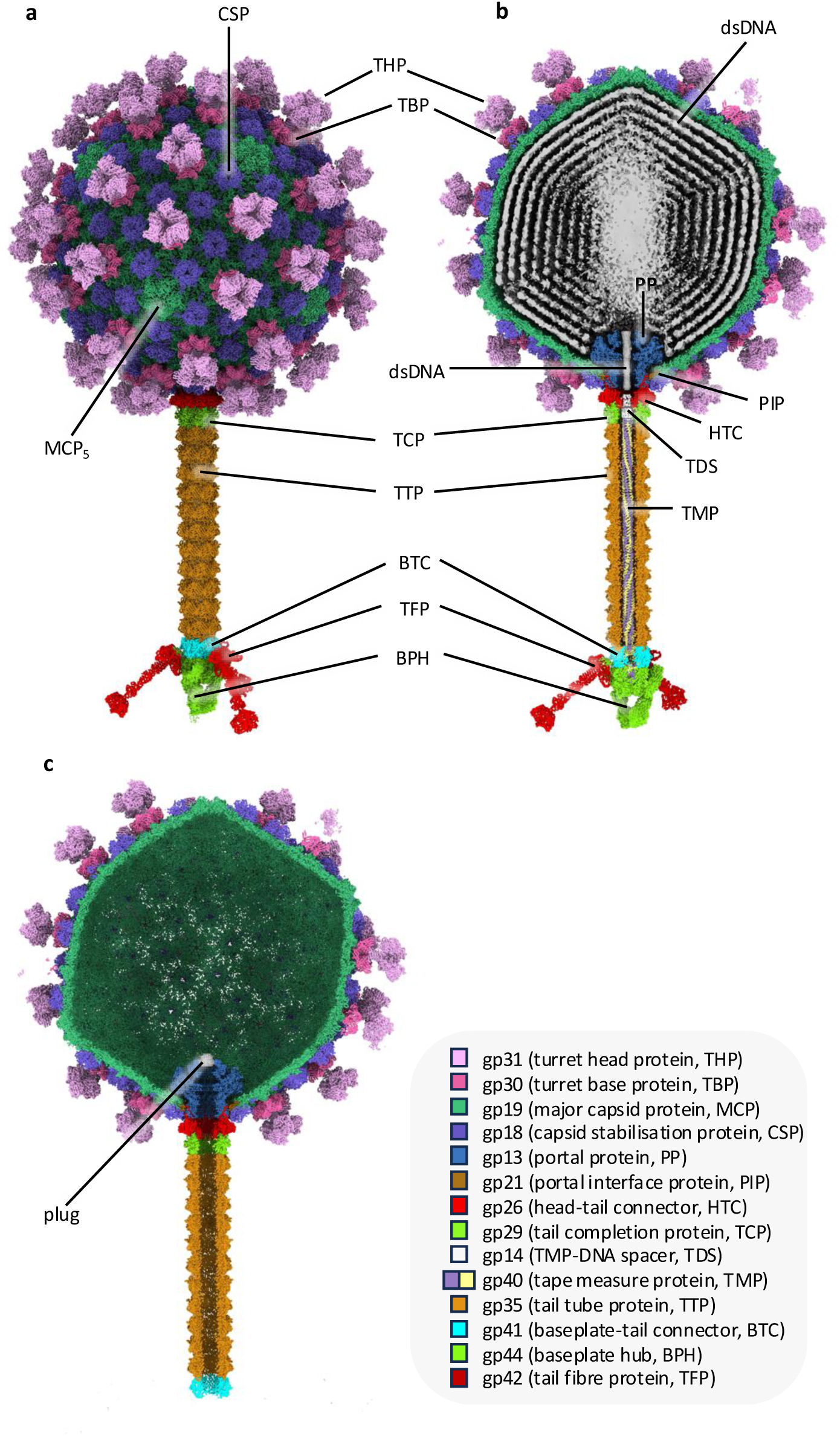
Overall structure of HFTV1. **a,b,** structure of head-tailed and turreted archaeal virus HFTV1 virion containing dsDNA in side view (**a**) and cross-section (**b**). **c**, cross-section of the empty virus, where dsDNA, tape measure protein, baseplate hub and tail fibres are missing. Instead, a plug-like density appears to cap the portal protein. The empty structure corresponds to a post-infection state.

Comparing the structures of the DNA-filled and empty virus (Fig. 1b,c) revealed that the latter lacked the baseplate and tail fibres, suggesting that these parts are jettisoned during DNA ejection. However, the length of the tail tube remains unaltered, demonstrating that it is rather stiff and does not contract during DNA ejection, as is typical for siphoviruses that infect bacteria [41,42]. Moreover, the heads of filled and empty viruses showed no significant differences, indicating that DNA release does not lead to any structural changes in the head.

Structural interpretation and atomic model building were informed by a proteomic analysis (see Materials and Methods), as well as AlphaFold2 [43] predictions (Supplementary Dataset 1). Mass spectrometry analysis of the infectious virions was performed, which led to the identification of several structural proteins (Supplementary Table 1). Only one protein band showed an exact match to the predicted amino acid sequence of the protein gp29 (GenBank accession QAS68862.1). gp21 and gp14 were not detected, but the hypothetical proteins gp20, gp17, and gp43 were identified, suggesting a potential role as structural proteins. Full peptide data are provided in in Supplementary Dataset 2.

AlphaFold2 was used to predict the structures of each of the 68 proteins encoded by the genome of HFTV1 [15], as monomers, and various multimers where hardware capacity allowed. This expedited the assignment and refinement of confidently annotated sequences and allowed candidate proteins with no identified proteomic counterpart to be shortlisted.

### The head has right-handed *T* = 7 icosahedral symmetry and is studded with turrets

Icosahedral symmetric averaging of the viral capsid produced a 2.36 Å map (Cryo-EM Flowchart 1; Supplementary Figure 11), which was used to build an atomic model. The capsid forms a right-handed (*dextro*) icosahedron of a triangulation number *T* = 7 (*h* = 2, *k* = 1; [44,45]), made up of 60 copies of the 7-protein asymmetric repeating unit (Figs. 1 & 2). While *T* = 7 icosahedral quasi-symmetry is characteristic of several bacterial siphoviruses [46,47], the handedness of HFTV1’s capsid is unusual, as most resolved structures of both tailed and untailed *T* = 7 icosahedral viral capsids display left-handed (*laevo*) symmetry [24,48–53]. To date, only a handful of other viruses have been shown to form *T* = 7 *dextro* capsids via structural analysis, including three tailed bacteriophages [54–56] and the untailed eukaryotic papovaviruses [57].

**Figure 2.**
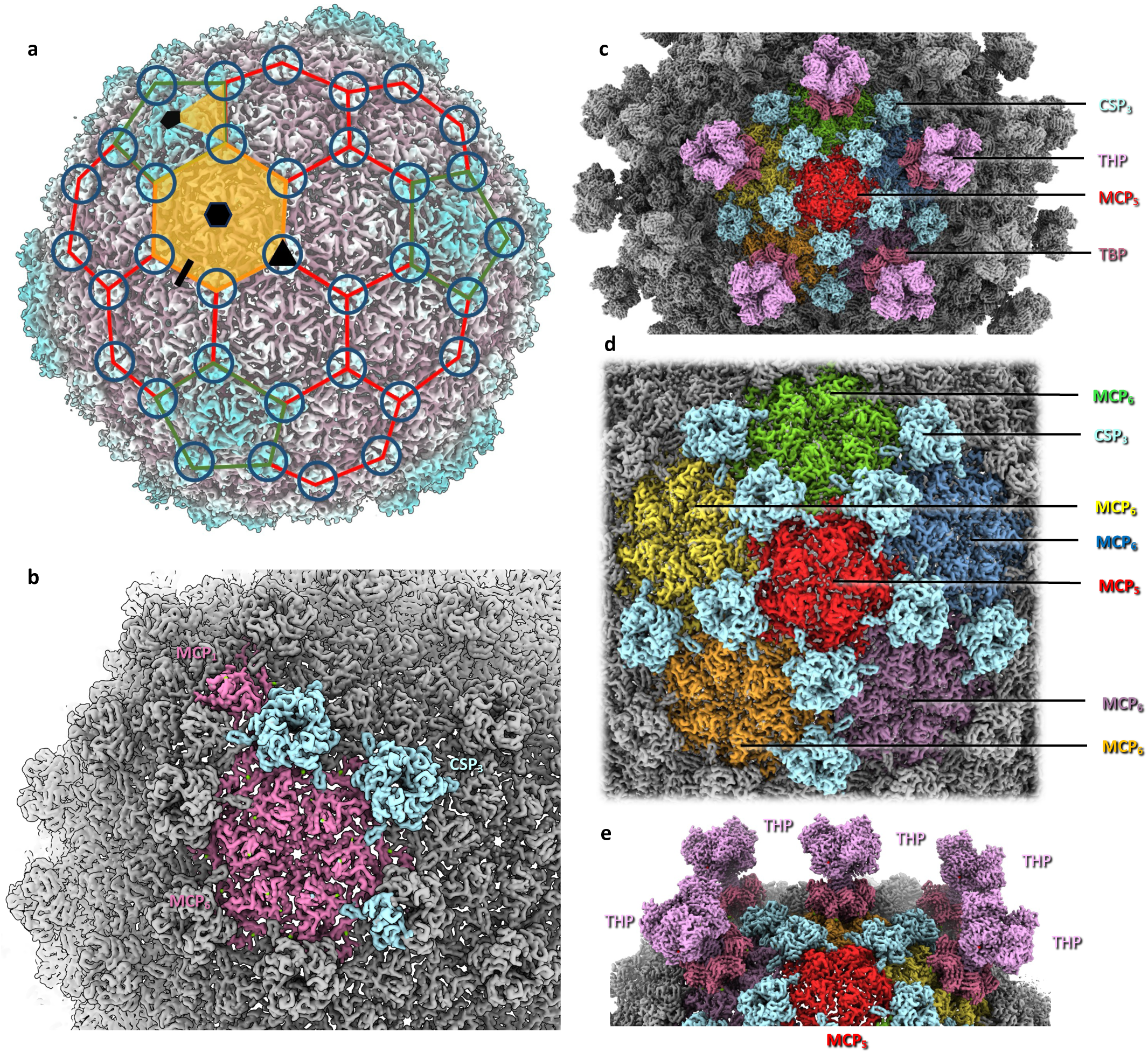
Structure of the icosahedral head of HFTV1. **a**, icosahedral capsid with major capsid protein hexamers (MCP_6_) outlined in red and major capsid protein pentamers (MCP_5_) outlined in green. The asymmetric subunit is highlighted in orange. Hexameric, pentameric, trimeric, and dimeric interfaces between MCPs are shown as black hexagon, pentagon, triangle, and line, respectively. The black pentagon, triangle, and line also demarcate 5-fold, 3-fold, and 2-fold axes of icosahedral symmetry, respectively. The location of capsid stabilization proteins (CSP) trimers is indicated by circles. The background map is coloured by radius in ChimeraX. **b**, close-up of the asymmetric subunit. MCP in pink, CSP in light blue. In **a** and **b** turrets have been omitted for simplicity. **c**, cropped view of the head, centred on an MCP_5_. The locations of the CSPs, turrets consisting of the turret base protein (TBP) and turret head protein (THP) are indicated. **d**, close-up of a central MCP_5_, surrounded by 5 MCP_6_ and 10 CSP trimers (CSP_3_). Turrets have been omitted for clarity. **e**, diagonal view of MCP_5_, showing that each MCP_6_ carries a turret, while all MCP_5_ remain unoccupied.

The base layer of the capsid consists of gp19, which is the major capsid protein (MCP). The HFTV1 MCP forms alternating pentamers and hexamers, which interact through asymmetric MCP trimers (Fig. 2a). In each trimer, the MCPs adopt three distinct conformations. Around the portal, the conformation of MCP is again different from those found in the asymmetric trimers, to accommodate the capsid-portal interface. Across the capsid, 13 different MCP conformations were found (Supplementary Figure 3a, Supplementary Movie 2).

The DALI server [58] revealed that HFTV1 MCP is structurally homologous to MCPs of tailed bacteriophages. The top hits were MCPs of Mycobacterium phage Che8 (PDB-8E16; [59]), Klebsiella phage Kp9 (PDB-7Y23), Mycobacterium phage Ogopogo (PDB-8ECN; [59]), and bacteriophage HRP29 (PDB-8ELD). Interestingly, the MCP of HFTV1 is missing the N-terminal amino acids 1–100 compared to the protein sequence predicted from the genome (Supplementary Figure 3b), suggesting that the protein is post-translationally processed, possibly during the maturation of the virus. Lacking the first 100 amino acids, HFTV1 MCP fits the classical HK97 fold. The interior-facing surfaces of the MCP are highly negatively charged, which has been suggested to prevent the packaged DNA from sticking to the capsid, thus facilitating rapid release upon infection [60].

The trimeric MCP interfaces are bound by trimers of gp18, the capsid stabilisation protein (CSP; Fig. 2). Across the icosahedral capsid, these CSP trimers are arranged in a series of hexagonal and pentagonal rings, which coincide with the faces and vertices of the icosahedron respectively (Fig. 1a, 2). HFTV1 CSP binds the outer surface of each MCP trimer, thus stabilising the capsid architecture.

According to DALI [58], HFTV1 CSP is homologous to capsid proteins from tailed bacteriophages, such as the head fibre gp8.5 N base of bacteriophage phi29 (PDB-6QYY; [61]), the capsid stabilising protein of the marine siphovirus TW1 (PDB-5WK1; [53]), and the Cyanophage Pam3 capsid protein (PDB-8HDT; [62]).

In the MCP hexamer, each subunit contains two intermolecular Mg^2+^ ions. Moreover, the interfaces between two MCPs within the hexamer coordinate two additional Mg^2+^ ions each. Another Mg^2+^ ion is coordinated at the MCP-CSP interface (Supplementary Figure 4a). In the MCP pentamer, each subunit coordinates three intermolecular Mg^2+^ ions instead of two. One Mg^2+^ ion is located at each MCP-MCP interface and one Mg^2+^ ion at the MCP-CSP interface. Additionally, each CSP trimer binds a potassium ion on a molecular three-fold axis (Supplementary Figure 4b). The extensive intra- and intermolecular metal ion coordination suggests a stabilising role of these divalent ions.

### The DNA packaging in the head is revealed

Aiming to understand how the dsDNA genome packs into the capsid of HFTV1, we first removed the capsid protein density from an unsymmetrised (C1) map of the capsid via particle subtraction and then refined the remaining DNA density using CryoSPARC [63] (Cryo-EM Flowchart 2; Supplementary Figure 14; Fig. 3a). To improve the resolution of the innermost layers, as well as the poorly resolved areas close to the portal and at the opposite pole of the capsid, we performed particle subtraction of these areas and refined them separately (Cryo-EM Flowchart 2; Fig. 3b). We found that the DNA is arranged in ten concentric shells (Fig. 3b). Two termini of the dsDNA were observed; one projecting through the portal (Fig. 3a) and the other likely located in the outermost layer, tracing the circumference of the portal (Fig. 3d).

**Figure 3.**
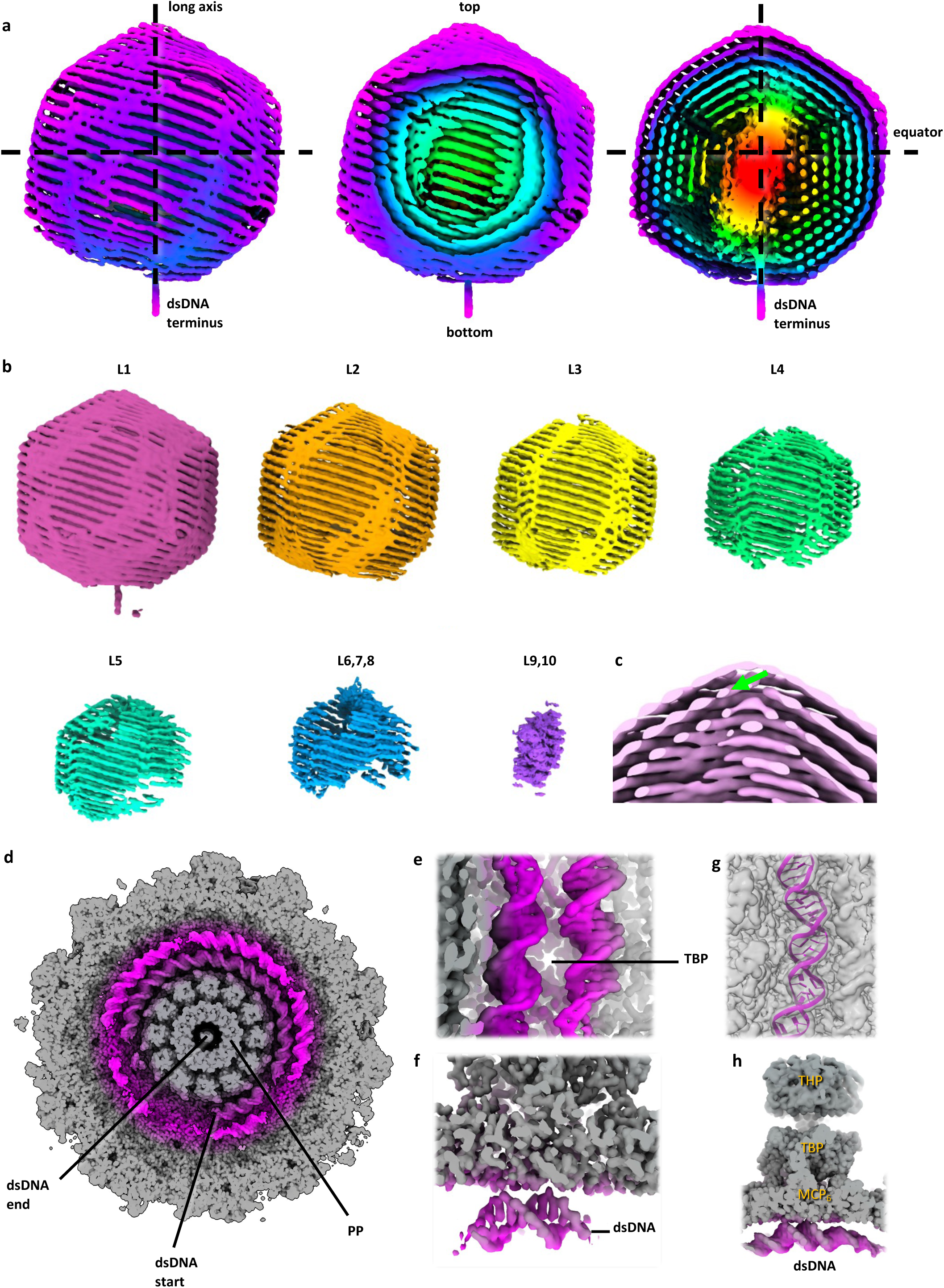
dsDNA spooling inside the virion head. **a**, cryo-EM map of the 10-layered dsDNA spool in full view (left), with the first 4 layers cropped (middle), and cropped through the centre of the dsDNA (right). The dsDNA emerges from the bottom of the spool at the portal. Note that the spooling direction of the DNA is inclined with respect to the equatorial plane of the capsid. **b**, the layers (L) of the dsDNA shown separately. The outer layers L1-L3 are resolved best, while layers L4-L10 show an increasing proportion of unresolved areas, indicating disorder. **c**, close-up of the spool at the opposite end to the portal. The green arrowhead indicates dsDNA descending from layer 1 into layer 2. **d**, view of the portal area of the capsid (PP, portal protein). Layer 1 of the dsDNA (magenta) is particularly well resolved around the portal and the twist is clearly visible. The putative terminus of the DNA molecule is indicated. **e-h**, various views of two parallel DNA strands of Layer 1 beneath a MCP hexamer (h). In this region, base pairs are resolved (**e,f**) allowing a section of dsDNA to be modelled (**g**).

In maps of the capsid-tail interface where no symmetry was applied, the DNA surrounding the portal is better resolved than in other areas of the capsid. This indicates that the dsDNA is well ordered and aligned with the capsid, suggesting that it is perhaps anchored in this region (Fig. 3d–h). The twist and base pairs of the dsDNA can be clearly discerned (Fig. 3d–f) and modelling a dsDNA molecule into the density indicated that it has B-DNA conformation (Fig. 3g), as opposed to the A-conformation reported for Sulfolobus islandicus rod-shaped virus 2 (SIRV2) and several other viruses infecting hyperthermophilic archaea [64–67]. In other dsDNA viruses such as siphoviruses and herpesviruses, tightly packed layers of DNA are observed, with higher resolution in the outer layers, as well as around the portal [68,69] and B-form DNA is also commonly observed [68,70,71].

Interestingly, the dsDNA is particularly well resolved at hexagonal capsid faces that coincide with the turrets proximal to the portal (Fig. 3e–h). Here, two adjacent DNA strands of the first layer align with the interior-facing side of the MCP (Fig. 3e–h). It is conceivable that the hexagonal vertices act as anchoring points for the DNA and help coordinate the packing.

The very centre of the capsid shows greater density than the outermost DNA layers. To investigate whether this central density consisted of a specialised region, potentially housing cargo proteins, we attempted masked refinement of this area. The resulting map (Fig. 3b, Layers 9 and 10) indicated that the central density mostly consists of DNA, namely the two central layers that are more densely packed than the ten outer layers surrounding them.

From the terminus near the portal, the DNA forms spooled layers with an incline of 17° relative to the base of the portal. The centre-centre distances between the DNA strands measure 25 Å within the same layer, as well as across layers 1–7. This agrees with previous observations of the spooled organisation of dsRNA, in which strands in the same layer are separated by 26 Å [72]. However, the interstrand distance across dsRNA layers is 29 Å [72], indicating that the dsDNA within HFTV1 displays tighter packing. The dsDNA also displays variations in the inclination across layers, with the outer six layers (1–6) displaying similar inclinations of 16°, 18°, 17°, 17°, 17°, and 16°. Layers 7–9 were not resolved well enough to accurately measure the incline of the DNA.

In the inner core, the distances between DNA strands are reduced to 23 Å and the incline is 11° (Fig. 3a,b). This contrasts with previous observations of dsRNA, in which the inner layer does not follow the spooled organisation and instead forms a barrel-like structure [72], whereas in the case of HFTV1, it appears that the dsDNA maintains the spooling organisation throughout. Furthermore, dsRNA structures display multiple points where the strands move between layers, corresponding to the capsid vertices [72]. In contrast, the dsDNA of HFTV1 forms depressions in these regions but does not seem to move between layers (Fig. 3a). It is currently unclear what causes these depressions, as no protein or other densities were observed that could be responsible.

Our data also did not conclusively reveal any protein densities that could facilitate this closer packing in the centre, so the mechanism for the observed packaging transition currently remains enigmatic. As we were not able to clearly resolve the very centre of the core, the presence of auxiliary packaging proteins remains a possibility. Analysing the genome of HFTV1 using InterPro [73], HHpred [74], and arCOG [75], we identified various gene products with predicted DNA-binding functions, including various Zn-finger proteins (Supplementary Datasets 3 and 4), however, the precise function of these proteins remains to be elucidated by further experimentation.

While the DNA could be traced for large areas, pockets of disorder remain. This particularly applies to the poles of the spooled dsDNA. Due to the incline of the spooled DNA, these regions do not coincide with the poles of the capsid, but instead with triangular faces near the poles. The top pole of the dsDNA also shows another degree of disorder, likely due to different paths of descent into the inner layers (Fig. 3c), which have been averaged together and could not be separated in classification.

Based on our data, we can describe the spooling path of the dsDNA in HFTV1. From the well-ordered terminus circling the portal at the bottom pole, the DNA winds up towards the top pole, forming the outer layer. At the pole opposite the portal, the DNA then descends into the second layer (Fig. 3c). From here, the DNA likely spools back down towards the pole at the portal, where the spooling direction is reversed again. This spooling path repeats nine times, until the DNA finally reaches the central core and protrudes back through the portal.

### The turrets have a putative catalytic deacetylase domain

Each of the hexagonal MCP of HFTV1 forms the base of a turret. In icosahedral and unsymmetrised averages of the capsid, the turrets were poorly resolved – an indication for flexibility. To improve the resolution of the turret map, localised reconstruction techniques were employed to take advantage of both the symmetric distribution and local symmetry of the turrets (Cryo-EM Flowchart 1). Particles were symmetry-expanded around the five-fold-symmetric head-tail axis, sampling the five particularly well-defined turret complexes immediately adjacent to the portal. Three-fold rotational (C3) averaging focused on a single turret position produced a 2.36 Å resolution map (Cryo-EM Flowchart 1). Turrets were found to consist of two separate proteins; a base formed of a hexamer of gp30 (turret base protein, HFTV1 TBP; Fig. 4), and a trimeric turret head protein gp31 (HFTV1 THP; Fig. 4). Extensive Mg^2+^ ion coordination involving 12 ions per turret (2 per TBP subunit) stabilises the interaction between TBP and MCP (Fig. 4f, Supplementary Figure 4a).

**Figure 4.**
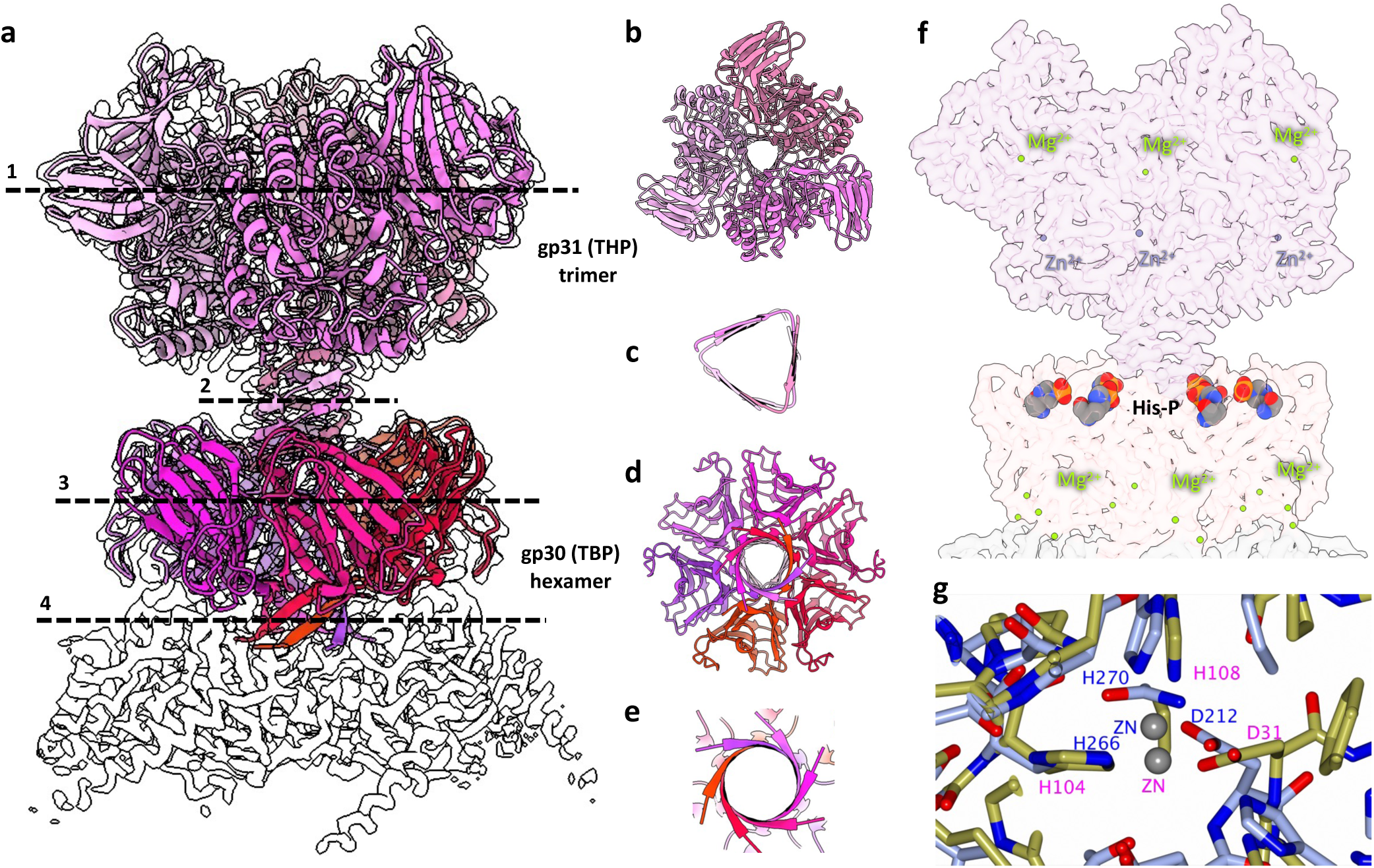
Structure of the turret. **a,** side view of the turret, consisting of a turret head protein (THP) trimer and a turret base protein (TBP) hexamer. Cross-sections through THP (1), the THP stalk (2), the TBP (3), and the TBP stalk (4) are indicated. The underlying MCP hexamer is shown in white. **b-e**, 90° rotated cross-sections corresponding to 1-4 in panel a, respectively. **f**, transparent map of the turret (side view) with coordinated Mg^2+^ (green) and Zn^2+^ (purple), as well as phosphohistidine residues (multicolour) indicated. **g**, superposition between the proposed active site of HFTV1 THP (blue) and that of the polysaccharide deacetylase from *Mycolicibacterium smegmatis MC2 155 (*PDB*-*3RXZ; yellow).

Both the TBP hexamer and THP trimer have mushroom-like architectures. TBP inserts into the MCP trimer with a short stalk-like protrusion formed by a 6-stranded β-barrel (Fig. 4e). The THP trimer then inserts into the TBP hexamer with a longer, 12-stranded barrel with a triangular cross-section (Fig. 4c), elevating the THP above the TBP. Six phosphohistidine residues are found at the hexamer-trimer interface between the TBP and THP (Fig. 4f). While all these modified histidines belong to the TBP, three of them directly interact with the N-termini of the THP trimer via salt bridges. These salt bridges contribute to the stability of the interaction between the THP and TBP.

Alongside three Mg^2+^ ions, the THP trimer includes three catalytic Zn^2+^ coordination sites (Fig. 4f,g, Supplementary Figure 4d). Analysing the structure of the THP using DALI [58] and NCBI Basic Local Alignment Search Tool (BLAST) [76] indicated a similarity between THP and polysaccharide deacetylases from multiple species (Supplementary Dataset 5). Superimposing the structure of the THP with that of the polysaccharide deacetylase from *Mycobacterium smegmatis* (PDB-3RXZ) results in a close match between the catalytic domains of both proteins (Supplementary Figure 5).

InterProScan [77] reports that the deacetylase domain encompasses residues 199–413, ending at the C-terminus of the THP. However, a second conserved domain is also reported, corresponding to a galactose-binding domain-like superfamily (G3DSA:2.60.120.260), which spans residues 57–198 and makes up the second externally facing subunit of the THP. Notably, following model building, the sequence of gp30 needs to be reannotated as the density contained more amino acids than suggested by the annotated genome (Supplementary Figure 6).

Interestingly, we found that the turrets are assembled in two possible orientations, where the turret head trimer can be rotated by 60 degrees with respect to the turret base hexamer. This rotational flexibility may increase the chance of binding between turrets and the S-layer of *H. gibbonsii*.

### The portal and its integration into the capsid

The portal complex integrates into one of the pentagonal MCP facets of the capsid. Particles recentred on the capsid-tail interface underwent 12-fold rotational averaging to produce a 2.34 Å resolution map of the portal protein (Fig. 5a; Cryo-EM Flowchart 1). The portal consists of a dodecamer of gp13 (portal protein, PP; Fig. 5a,c), which spans the capsid through one pentagonal vertex (Fig. 5b), followed by a dodecamer of gp26 (head-tail connector, HTC) and a gp29 hexamer (tail completion protein, TCP; Fig. 5a). The symmetry mismatch between the MCP pentamer and the PP dodecamer is interfaced by the intermediary portal interface protein gp21 (PIP; Fig. 5a,b). Modelling the PIP into the map showed that its annotated genome sequence was eight residues shorter than the modelled one, suggesting that the starting methionine is different from that originally annotated (Supplementary Figure 7). At the portal-capsid interface, one of the MCP-bound magnesium ions is replaced by an arginine of the PIP. In addition, the PIP also has a phosphohistidine residue at position 17, which forms an H-bond with neighbouring MCP monomer (Fig. 5f,g).

**Figure 5.**
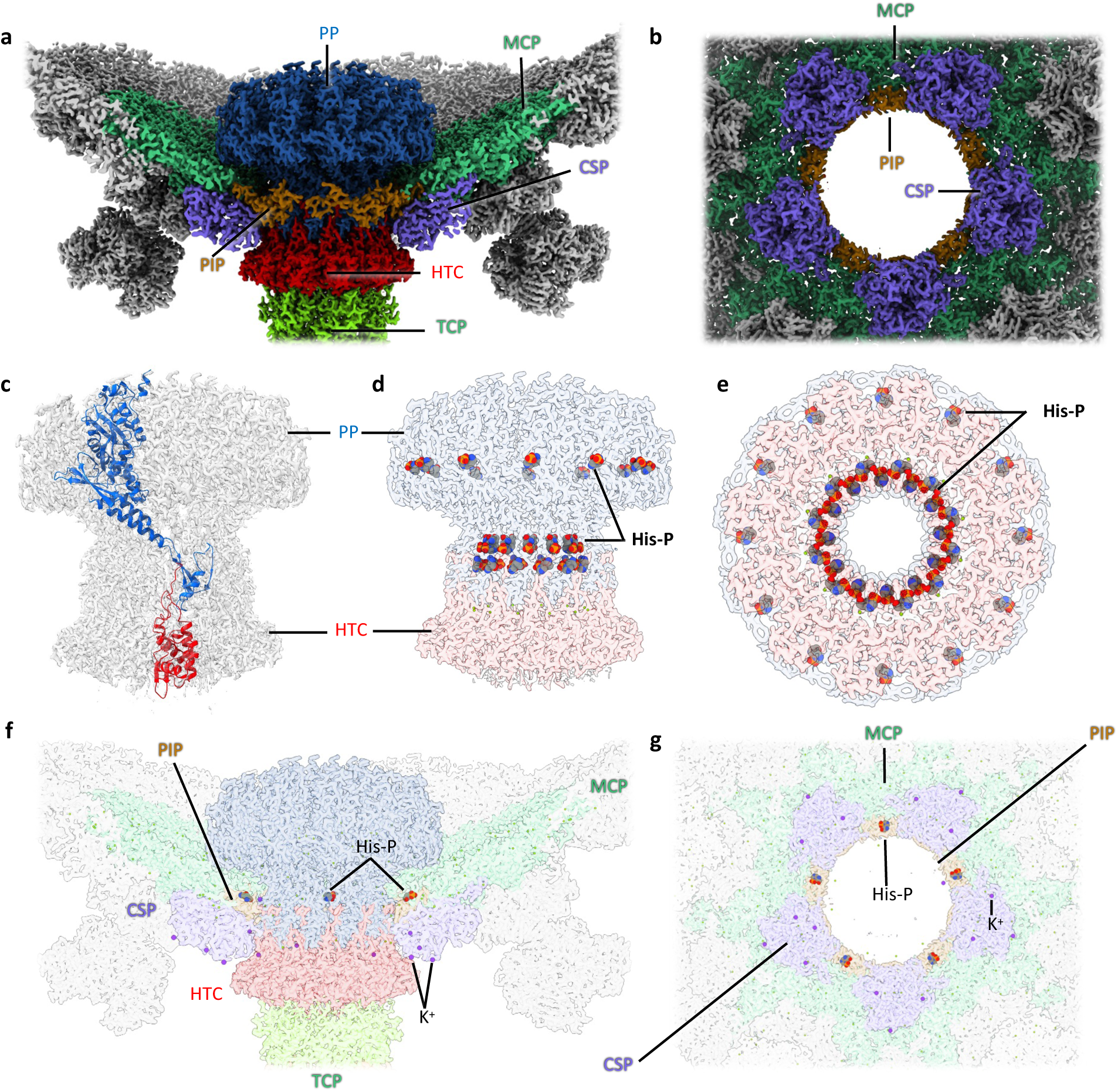
The portal. **a**, cross-section through the portal region with the portal protein (PP, gp13, blue) in the centre. A pentameric ring of gp21 (portal interface protein, PIP, brown) integrates the portal 12-mer into the 5-fold vertex of the capsid. **b**, top view of the portal interface, the PP has been omitted for simplicity. **c**, cryoEM map of the portal in transparent grey, with PP (blue) and head-tail connector (HTC, gp26, red) monomers shown as atomic models in ribbon representation. **d,e,** cryo-EM map of the portal in side view (**d**) and top view (**e**). PP in transparent blue and HTC in transparent red. The PP 12-mer contains 36 phosphohistidine residues (His-P; red and blue spheres). 24 of this form a ring lining the portal tunnel. **f,g**, transparent views of **a** and **b**, respectively. Phosphohistidine in the portal-capsid interface, as well as coordinated K^+^ (purple dots) are indicated.

In the mature virion, the terminus of the dsDNA protrudes through the central vestibule of the PP (Fig. 6b,c), which is lined with a ring of phosphohistidine residues (Fig. 5d,e). These phosphohistidines add negative charge to the portal tunnel and likely disfavour any DNA binding to the portal, thus facilitating rapid DNA release during infection. Comparing the portal structures of the DNA-filled and empty capsid shows that the latter has a plug-like density, which is missing in the former (Fig. 6b,c). The structure of the PP itself does not show any significant differences between the two states, indicating that conformational changes in this protein are not required to facilitate DNA release during infection.

**Figure 6.**
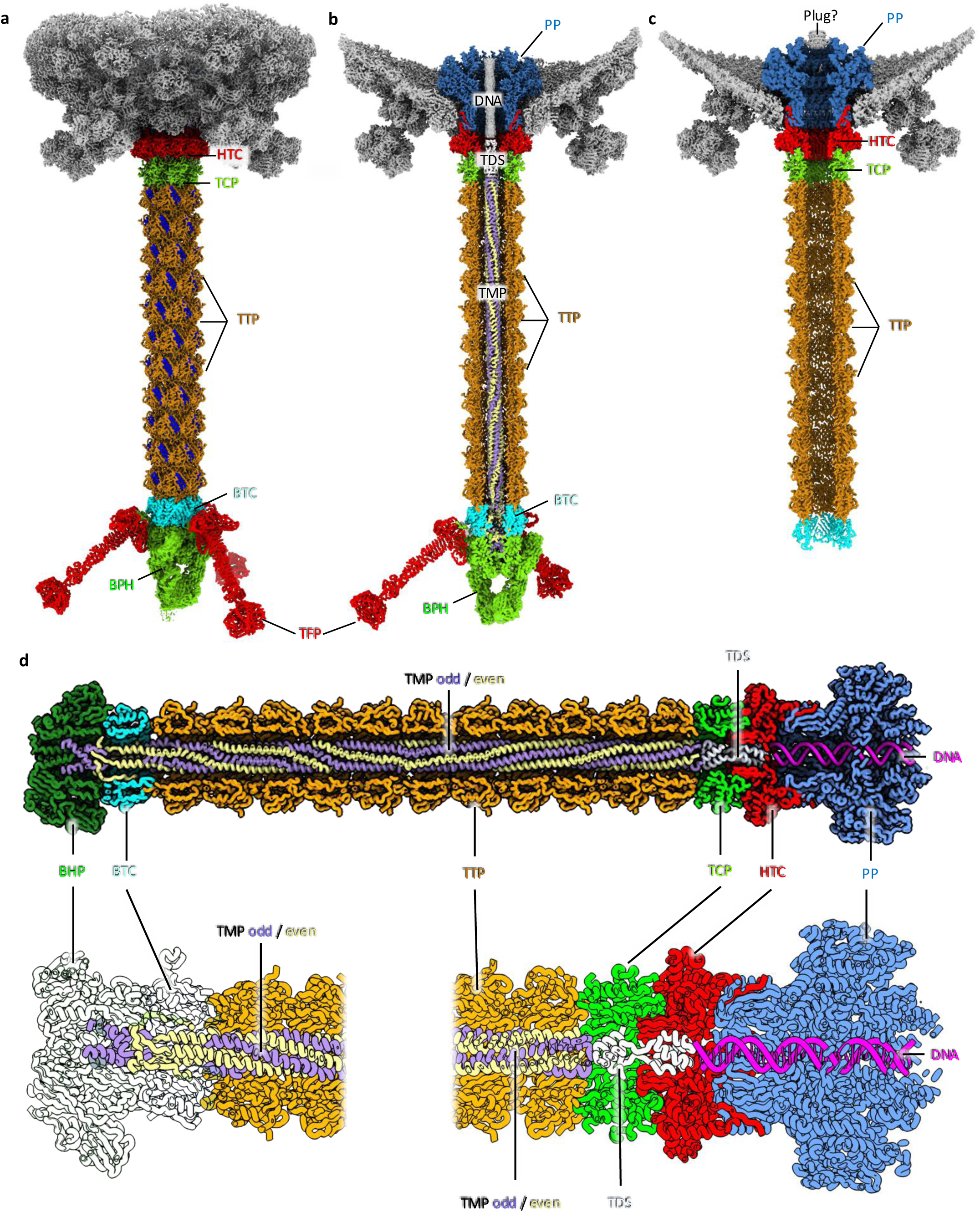
The tail. **a,** full structure of the tail consisting of a dodecameric head tail connector (HTC), a hexameric tail completion protein (TCP), 11 tail tube (TTP) hexamers, a trimeric baseplate tail connector (BTC), a trimeric baseplate hub (BPH), and 3 trimeric tail fibres (TFs). The same pair of β-strands per TTP monomer is highlighted in blue, showing the 23.6° rotational offset between the TTP hexamers. **b**, cross-section of the tail tube, revealing a hexameric tape measure protein (TMP), projecting from a cavity in the BPH to a vestibule formed at the TCP. The TMP and dsDNA are separated by a spacer (TDS), and the dsDNA traverses through the portal into the capsid. **c**, structure of the tail of the empty virus. The dsDNA, TMP, and baseplate are missing. Instead, a density appears to plug the capsid-facing side of the portal. Otherwise, the portal and the components of the tail tube are unchanged in width, length, and conformation, indicating a rigid structure. **d**, close-ups of cross-sections through the atomic model of the DNA-loaded tail.

### A complete structure of the tail and tape measure protein

Particles recentred on the tail underwent refinement applying C3 symmetry, which resolved the tail at 2.45 Å resolution (Fig. 6a,b). The tail region begins with dodecameric gp26 (head-tail connector, HFTV1 HTC), which interacts with the portal protein to maintain a conduit through which DNA can pass. This is followed by hexameric gp29 (tail completion protein, HFTV1 TCP), and subsequently, 11 copies of hexameric gp35 (tail tube protein, HFTV1 TTP) making up the bulk of the tail (Fig. 6a,b,d). Each hexameric segment of the tail is rotationally offset with respect to the next by 23.6° (Fig. 6a).

The tail forms a hollow tunnel of 36 Å in diameter. In the DNA-filled virus, this tunnel houses gp40 (the tape measure protein, HFTV1 TMP), a hexameric coiled-coil structure with alternating C-terminal conformations. The TMP reaches from the central vestibule of the first TTP near the portal to a central cavity in gp44 (baseplate hub, HFTV1 BPH) (Fig. 6b,d). Based on our map, we could build the tape measure protein (TMP) as a full atomic model, with all side chains present. The C-terminus of TMP (residues 317– 341) is well-defined in the density, with odd-numbered subunits forming α-helical structures docked into the baseplate hub (Fig. 6d). The even-numbered subunits, on the other hand, form a β-sheet structure folding back toward the N-terminus (Fig. 6d). Residues 2–316 fold into α-helical structures with three-fold symmetry, spanning most of the tail length. In the extended N-terminal part of the protein, the density remained ambiguous in various regions, and side chains were poorly resolved.

The density within the portal protein clearly corresponds to dsDNA (Fig. 6b). Additionally, we observe two densities between the DNA and the TMP, located within the TCP and HCP proteins (Fig. 6b). Using AlphaFold3 [78], we predicted the structures of oligomers for all short HFTV1 and proteins of unknown function, resulting in a dimer of gp14 as the best match for this density. The gp14 dimer forms two globular domains, one at the N, and the other at the C-terminus, connected by short loop regions (Fig. 6d). Due to the strong features of three-fold symmetry of the TMP, we were unable to align particles based on the weaker features of gp14 and the neighbouring DNA, leading to the averaging out of secondary structure details of this region. However, superimposing the dimer of gp14 as a rigid body resulted in a reasonable fit. Due to its location between the TMP and DNA, we name this protein the TMP-DNA spacer (HFTV1 TDS). In the empty virus, the TMP and TDS are absent, demonstrating that they are released during DNA ejection.

### The baseplate hub forms an asymmetric trimer carrying three tail fibres

The tail of 11 hexameric TTPs is followed by trimeric gp41 (baseplate-tail connector; BTC), which in turn interfaces with the trimeric BPH (Fig. 7a,b). Notably, the BTC protein is a pseudo-hexameric trimer. Each subunit consists of two homologous domains, which have likely arisen through a gene duplication and fusion event (Fig. 7c,d). The BPH complex connects to the BTC via a triangular ring, from which three globular domains project (Fig. 7e). The BPH coordinates Mg^2+^ and K^+^ ions that likely contribute to the stability of the complex. Curiously, these domains show an asymmetric configuration (Fig. 7e). While two of the domains are bound to each other, one remains unbound. As the binding interfaces of these three domains are equal, it can be assumed that they are free to interchange (Supplementary Movie 3). This suggests that the three BPH domains may undergo dynamic cycles of binding and unbinding, which in turn may be important for the ability of the virus to dock to or penetrate the S-layer.

**Figure 7.**
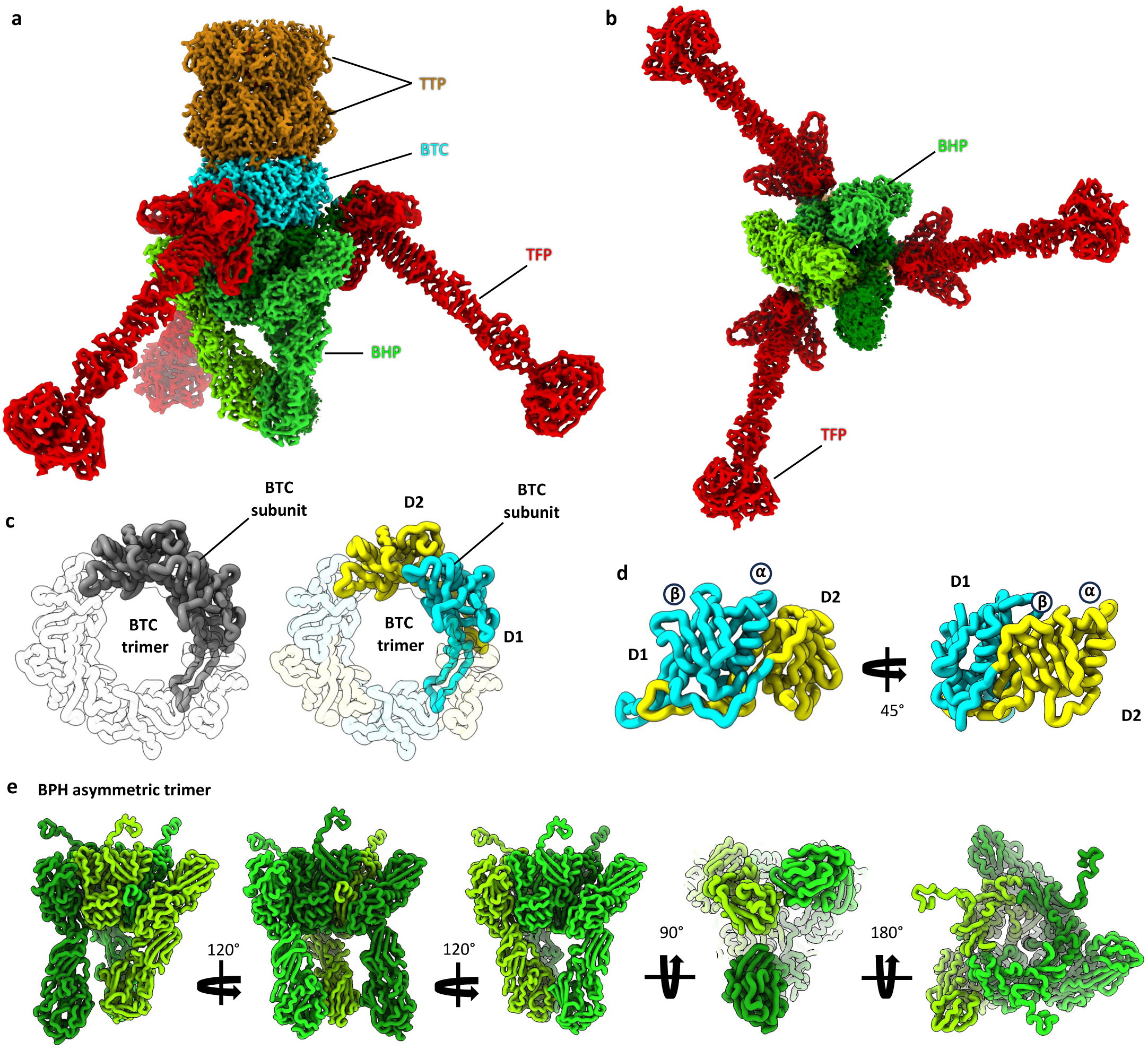
The baseplate. **a,b**, structure of the HFTV1 baseplate in side (**a**) and end-on view (**b**). **c,** left panel, BTC trimer in top view with one of the subunits coloured in grey and the other two transparent; right panel, the same subunit is coloured by domain (D1, cyan and D2, yellow). **d**, two side views of the BTC subunit, rotated by 45°, showing that both domains adopt the same fold. Both domains consist of a β-sheet followed by an ⍺-helix (annotated ⍺ and β). **e,** various views of the asymmetric baseplate hub protein (BPH) trimer (monomers coloured with different shades of green). Three extended domains project downwards from the core of the BPH. Each of the domains has a different conformation. While two interact directly, the third is unbound and thus more flexible.

The BPH is also the anchor point for three tail fibres (Fig. 7a,b). The tail fibres are anchored at the N-terminus of each BPH subunit and radiate away in a ∼45° angle with respect to the BPH. This interaction is stabilised by extensive hydrogen bonds and Mg^2+^ ion coordination. A DynaMight [79] analysis revealed that the tail fibres are highly flexible with respect to the BPH, likely aiding their ability to dock to the S-layer at the cell surface (Supplementary Movie 4).

Alphafold2 [43] suggested that the tail fibres are formed as a homotrimer of gp42 (tail fibre protein, HFTV1 TFP) and this was in agreement with atomic model building via ModelAngelo [40]. Three TFPs come together to form a base, followed by a β-helix, a stalk, and a head. The TFPs coordinate Mg^2+^ and Zn^2+^ ions (Supplementary Figure 4c), which likely stabilise the trimer interface. According to DALI, the BPH shows some homology with xylanases and glycosaminidases (Supplementary Dataset 5). For the TFs, DALI suggests some homology with the CspB protease from *Clostridium perfringens* (PDB-4I0W; [80]), albeit the Z-score is very low. This may hint at the BPH and TFs playing a role in binding to the surface glycans of *H. gibbonsii* and even the degradation of the S-layer. However, further experimentation will be required to confirm this supposition.

### Coordinated magnesium ions are crucial for virion stability

We asked if the multiple intra- and intermolecular Mg^2+^ ions coordinated throughout the baseplate hub, tail fibres, turrets, and head-tail connector proteins (Supplementary Figure 4a) have a stabilising role. To test this, we depleted Mg^2+^ from infectious virus particles using chelating agents. This resulted in a loss of infectivity (Supplementary Figure 8) and fragmentation of the virus particles (Supplementary Figure 9), showing that Mg^2+^ indeed aids the structural integrity of the virus. The extensive ion binding within the capsid is likely a result of HFTV1 evolving in a highly saline environment, where Mg^2+^ ions are abundant.

### Conservation and divergence of HFTV1 compared to archaeal and bacterial head-tailed viruses

Despite the fact that HFTV1 infects archaea, its overall architecture is strikingly reminiscent of bacterial siphoviruses, which are characterised by an icosahedral head and a non-contractile tail. To investigate whether HFTV1 is indeed related to bacteriophages, we performed a homology analysis for each component protein of HFTV1 using the DALI server [58]. Probable homology (Z-scores between 8 and 20) with proteins of bacteriophages was determined for the majority of HFTV1 structural proteins. These include (in descending order of Z-score) major capsid protein, capsid stabilisation protein, tail completion protein, tail tube protein, tail-baseplate connector, and tail fibre protein (Supplementary Dataset 5).

Interestingly, DALI suggested that the portal protein and the baseplate hub protein of HFTV1 are definite homologues of phage counterparts (with Z-scores of >20). For the portal and the baseplate hub, the greatest Z-scores were identified in Thermus phage G20c portal (PDB-4ZJN) and prophage MuSo2 43 kDa tail protein (PDB-3CDD), respectively.

Similarity, but no definite homology could be determined for the head-tail connector, and the portal interface protein PIP had no known homologous hits whatsoever, suggesting that this part of the capsid has a unique origin. Interestingly, the turret head protein also did not show any homology with bacteriophage proteins, but instead with bacterial polysaccharide deacetylases (top Z-score 18.6 with an enzyme from *Mycolicibacterium smegmatis* MC2 155 [PDB-3RXZ]). Another intriguing homologue was found in an Endo-1,4 β-xylanase from the termite species *Trinervitermes trinervoides* (PDB-7AX7; [81]) with a Z-score of 18. For the turret base protein, only one probable homologue could be identified, a phage-like element PsbX protein from *Bacillus subtilis* strain 168 (PDB-6IA5; [82]).

In conclusion, these data hint at the possibility that HFTV1 is directly evolutionarily related to tailed bacteriophages and that core structural proteins, particularly those forming the portal, capsid, tail, baseplate, and tail fibres, could be products of horizontal gene exchange between bacterial and archaeal viruses. Other proteins that convey specific adaptations to the archaeal host, such as the turret heads proteins, appear to have evolved high specificity for HFTV1.

## Discussion

Based on our structure of the HFTV1 virion, a comparative analysis of the HFTV1 structural proteins that showed the presence of glycan binding sites, and previous observations of particle attachment, a multistep adsorption mechanism of viral binding can be deduced (Fig. 8). We propose that HFTV1 initially adheres reversibly to host glycans via the N-terminal carbohydrate-binding module of the turret head protein, which forms the outermost region of the turrets. Concurrently, the C-terminal region of turret head protein, which possesses putative glycoside deacetylase/hydrolase activity, starts degrading the glycan matrix of the host cell. This enzymatic activity weakens the protective Surface layer lattice of *H. gibbonsii*, which causes the rigid structure to loosen and expose the underlying membrane. The gradual degradation of the host glycans enables the viral particle to bring its tail apparatus closer to the cell membrane and may expose a new binding site, facilitating baseplate anchoring and viral reorientation. We suggest that this crucial step is mediated by the carbohydrate/galactose-binding domain on the C-terminal end of the baseplate hub. The baseplate, which serves as an anchor point on the cell membrane, secures the viral particle firmly to the host cell. This attachment by the baseplate, which is irreversible in many other viruses [83–85], reorients the virus so that the tail axis is aligned perpendicularly to the cell surface. The baseplate likely interacts with either remaining glycan residues on the S-layer or lipid-glycan carriers embedded in the membrane. It is conceivable that the potentially dynamic baseplate structure serves as a lock-and-key mechanism to open or disrupt the S-layer proteins, creating a pathway for virus genome entry. Trimeric baseplates with carbohydrate-binding domains have also been reported for bacteriophages [86].

**Figure 8.**
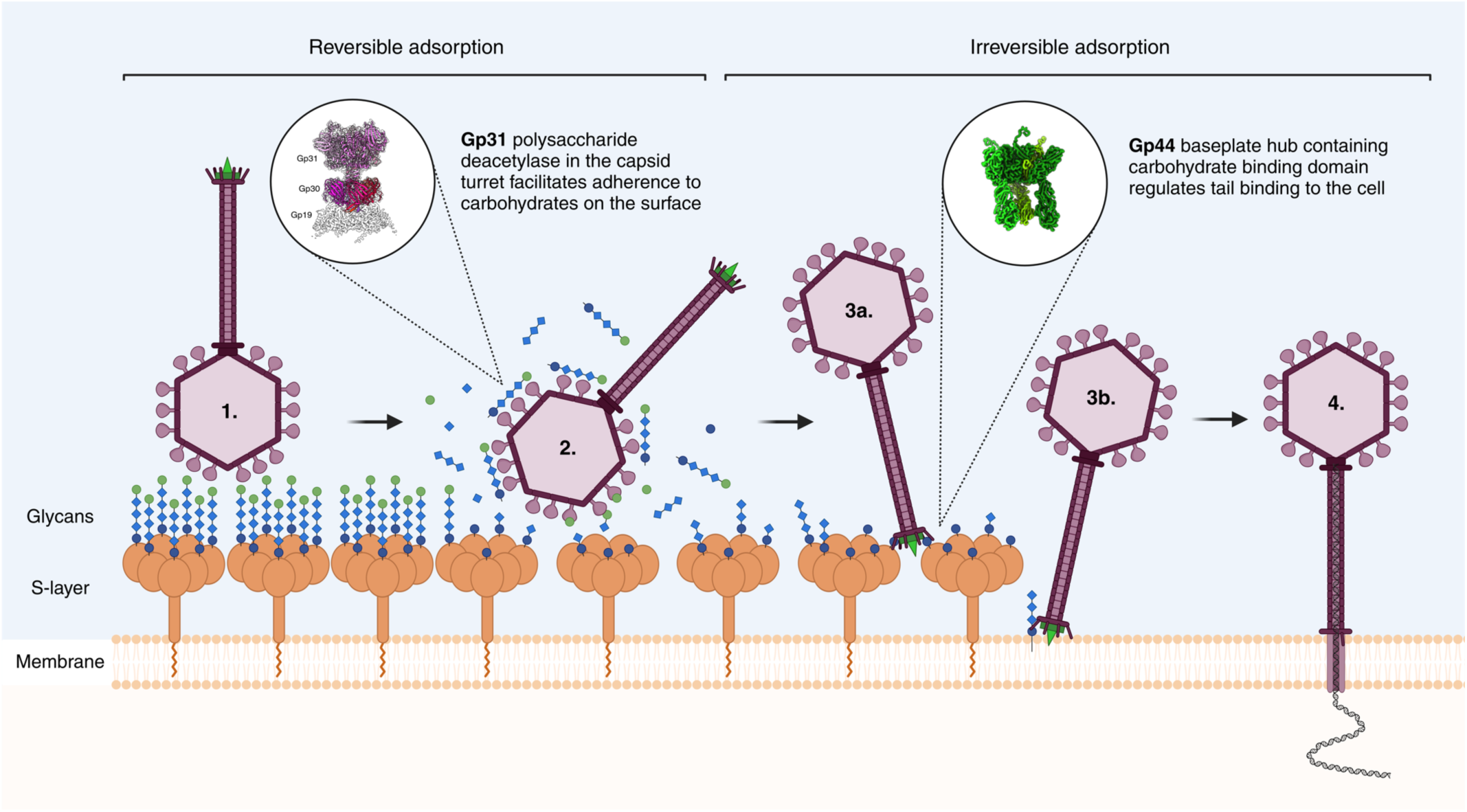
Proposed multistep receptor binding mechanism driving HFTV1 attachment to *Haloferax gibbonsii* LR2-5 and viral DNA delivery. **1.** HFTV1 binds reversibly to host glycans (chains of blue and green circles and diamonds) associated with the S-layer via the N-terminal domain of the THP (gp31). **2.** Simultaneously, the C-terminal polysaccharide deacetylase domain of the THP degrades the glycan matrix, weakening the S-layer of the host. **3a.** After glycan degradation, irreversible binding is mediated by a carbohydrate-binding domain on the BPH (gp44), which presumably binds to glycan residues on the S-layer. **3b.** Alternatively, it may bind to lipid-glycan carriers in the membrane. This secondary binding event reorients the particle. **4.** The BPH-mediated binding induces conformational changes that facilitate genome ejection into the host cytoplasm, accompanied by loss of the baseplate hub.

The tail has a length of 560 Å, which exceeds the distance between *H. gibbonsii* S-layer and membrane, which is ∼200 Å in *H. volcanii* [87–89] and likely similar in *H. gibbonsii*. This enables the virus to penetrate the S-layer and the membrane to create a conduit from the viral portal complex into the host cell, which is then used to inject the viral genome. Some phages have peptidoglycan hydrolysing activity in their tape measure proteins to degrade bacterial envelope components [28,90]. Proteolytic domains are not evident in the TMP of HFTV1, suggesting that the virus does not employ this proteolysis mechanism to overcome the S-layer of *H. gibbonsii*. Instead, the tail may penetrate the envelope mechanically, akin to a needle. Genome release is associated with structural changes in the distal tail region of HFTV1, including the detachment of the baseplate hub domain and dissociation of the tail fibres. These changes suggest that the baseplate and tail fibres are involved not only in anchoring the virus, but also in triggering the genome ejection process.

To our knowledge, this is the first example of archaeal virion containing structural proteins with carbohydrate-degrading domains. However, carbohydrate-binding and/or hydrolytic proteins have been abundantly reported as components of bacteriophages [91–94]. Several of these proteins are involved in hydrolysis of the bacterial peptidoglycan layer, presumably to gain access to the cell membrane. In the large majority of cases, these carbohydrate binding/hydrolytic proteins form the tail spike, baseplate, or are part of the tail fibres [91–94]. Interestingly, P22 podophage infecting *Salmonella typhimurium* has a peptidoglycan/murein hydrolase (gp4) inserted in the head structure that degrades peptidoglycan [95]. In addition, glycan binding domains have also been found in the capsid head of myophage T4, which are involved in adhering T4 to the mucus layer of the human gut, to increase the chances of encountering a susceptible *Escherichia coli* host [96]. Thus, multistep adsorption processes are common among bacteriophages (i.e. phage λ, T5, SPP1). Usually, binding to cell envelope saccharides is the first reversible step that maintains and transitorily orients the phage before specific receptor recognition occurs in the second step [97].

In HFTV1, the carbohydrate binding domain in the turret head protein likely serves as a first anchoring point of the viruses to the cell surface. This reversible binding step increases the chances that a successful irreversible binding (via the base plate and tail fibres) will take place, which then triggers DNA ejection and hence infection. As no homologues of the turret head protein were found in other haloarchaeal viral genomes, the described strategy could be specific to HFTV1. Indeed, HFTV1 was shown to have adsorption rate that is several orders of magnitudes faster than those known for other haloarchaeal viruses [35]. However, since quite a number of baTVs contain glycan binding proteins, and the archaeal cell surface is highly glycosylated, we expect that structural analysis of other haloarchaeal viruses will disclose the presence of similar glycan binding proteins in a variety of other haloarchaeal viruses.

The spooling of the dsDNA within the virus capsid is reminiscent of that displayed in several other dsDNA viruses of eukaryotes and bacteria [68,69]. The prevalence of this arrangement may arise due to this conformation being both electrostatically favourable as well as preventing entanglement upon injection into the host cell under extremely high pressure [33,68]. However, the axis of the spooling, which is out of alignment with the portal, is unique compared to other tailed dsDNA viruses, such as Epsilon15 [69], T7 [98], and herpes simplex virus (HSV) [68]. While HSV does display angled spooling (at least in the first layer), the axis of the spools still appears to align with the portal [68].

While it is possible that this conformation is unique to HFTV1, it is likely that within other dsDNA eukaryotic viruses and phages displaying linear spooling, such as HSV and Epsilon15, the averaging of different DNA conformations within the capsid may have caused the spools to appear linear, as mentioned in previous studies [99,100]. Since no studies of dsDNA organisation have been conducted in this much detail thus far, it is possible that the observed angled spooling is common across viruses, but further cryo-EM analysis of dsDNA viruses will be required to confirm this.

We observe areas of disorder within the spooled dsDNA, which is not unusual, especially for the innermost layers of viral genomes [33,68,98]. This disorder may be necessary to give the genome room to expand and shrink as mutations are introduced, and may also aid the unravelling of the spool as it is ejected. Interestingly, HFTV1 has an unusually dense packaging of central layers. The tight packing and bending at high angles displayed in this structure should be energetically unfavourable, given the negative charge and relatively rigid structure of dsDNA [101]. The curvature stress of the DNA may be enough to overcome the electrostatic repulsion, as demonstrated by continuum computations for the bacteriophage T7 [102].

During DNA packaging, as the internal pressure of the capsid increases, so do the durations of the burst and dwell phases of the packaging mechanism [103], resulting in an overall decrease of the packaging speed. Decreased packaging speed has been shown to increase disorder within the DNA layers [33,104]. This could suggest that disordered central layers may commonly be seen across other viruses with spooling conformations. However, in HFTV1 the central-most layers display a high degree of order, indicating the possible presence of histone-like proteins maintaining this structure. Future work aiming to increase the resolution of the packaged dsDNA may shed more light on the presence of such DNA-binding proteins in the capsids of archaeal viruses and beyond.

In summary, the structure of the tailed archaeal virus HFTV1 presented here provides structural evidence of the strong evolutionary relationship between tailed viruses of bacteria and archaea. As this is the first high-resolution structure of a tailed archaeal virus, it presents a major step forward in our understanding of archaeal viruses and opens up new research directions to answer fundamental questions in the field of virology.

Besides the overall structural similarity between bacterial and archaeal tailed viruses, HFTV1 shows unique features, such as the turrets that decorate the head and are likely responsible for glycan binding and degradation. Although the presence of a deacetylase on the capsid head is unique, a two-step binding mechanism relying on host glycan binding and degradation has also been observed for several baTVs and eukaryotic viruses. Thus, viruses from different domains of life seem to employ universal strategies for infection, relying on structurally distinct but functionally similar host-binding proteins on the viral capsid. The dsDNA spooling geometry revealed in the HFTV1 capsid exhibits previously unobserved characteristics, including a vertical spooling axis relative to the portal. We anticipate that future studies will uncover similar spooling arrangements as more high-resolution structures of DNA within capsids become available.

## Materials and Methods

*H. gibbonsii* LR2-5 cells and HFTV1 virions were cultured, and viruses were purified by precipitation and density ultracentrifugation in sucrose and CsCl as described previously [15,35]. Samples were concentrated by differential ultracentrifugation and resuspended in 18% SW buffer (2.47 M NaCl, 89 mM MgCl_2_, 85 mM MgSO_4_, 56 mM KCl, 3 mM CaCl_2_, 48 mM Tris-HCl, pH 7.2).

### Stability tests of the virions

The purified viruses were diluted 1:1000 in the following buffers: (i) 50 mM Tris-HCl, pH 7.2, (ii) 10 mM EDTA, 50 mM Tris-HCl, pH 7.2, (iii) 10 mM EGTA, 50 mM Tris-HCl, pH 7.2 and (iv) 18% SW (positive control), and incubated at 22°C. Number of infectious viruses were determined by plaque assay after 2 h and 24 h of incubation. To analyse the effect of magnesium depletion, the purified particles in 18% SW were collected (Beckman Coulter Airfuge rotor A95, 30 psi, 30 min, 22°C), resuspended in either (i) 200 mM EDTA, 50 mM Tris-HCl, pH 7.2 or (ii) 50 mM Tris-HCl, pH 7.2 (control) to obtain a particle concentration of 1 mg/ml, and analysed in a linear 10–40% (w/v) sucrose density gradient in 50 mM Tris-HCl, pH 7.2 (Thermo Scientific rotor TH641, 35,000 rpm, 1h 45 min (EDTA treated) / 35 min (control), 15°C). The gradients were analysed by absorbance profiling (A260 and A280) and collecting of fractions by Piston fractionator. Protein profiles were analysed by SDS-PAGE and infectivity of the light scattering bands were determined by plaque assay.

### Proteomics analyses by mass spectrometry

Purified HFTV1 virions were dissociated in SDS-PAGE loading buffer (4% SDS, 0.25M Tris, pH 6.8, 0.06% bromophenol blue, 0.5 mM dithiothreitol, 10% glycerol), boiled for 5 min at 95°C and the proteins were separated on 10–20% gradient polyacrylamide gels (Criterion precast gel, Biorad). The virus proteins were visualised by staining with Coomassie Brilliant Blue.

SDS-PAGE analysis of the highly infectious purified HFTV1 virions of high specific infectivity 2×10^13^ pfu/mg protein [35] revealed protein bands ranging in size from 250 to 20 kDa (Supplementary Figure 10). The proteins in the most prominent bands (bands 1 to 8, Supplementary Figure 10) were subjected to mass spectrometric analysis as described previously [105] at the Interfaculty Mass Spectrometry Core Facility (IMSC), University of Groningen. Briefly, in-gel digestion was performed on the excised gel bands with 150 ng trypsin. LC-MS based proteomics analyses were performed using 2/3 of these digests. LC-MS raw data were processed with Spectronaut (version 17.0.221202) (Biognosys) with a user-defined database containing 68 proteins (HFTV1 proteome, downloaded from GenBank, accession number NC_062739.1) using the standard settings of the directDIA workflow except that quantification was performed on MS1. For the quantification, the Q-value filtering was set to the classic setting and no imputation or normalisation was applied.

### Single particle data collection

Purified virions (1.4×10^14^ pfu/mL) were resuspended in 18% SW buffer and loaded onto R2/2 QUANTIFOIL 200mesh copper-carbon cryo-EM grids with graphene oxide support film. Grids were blotted using an FEI Vitrobot with force −1 for 5–6 seconds, under environmental conditions of 4°C, 100% humidity. Grids were plunge-frozen in liquid ethane, then transferred to and stored under liquid nitrogen.

Data were collected in two separate sessions at the Diamond Light Source Electron Bioimaging Centre (eBIC), utilising 300 kV Thermo Scientific Titan Krios TEMs with Gatan K3 and Falcon 4i direct electron detectors respectively. EPU software (Thermo Scientific) was used for automated data acquisition. The first dataset (Gatan K3) consisted of 10,236 52-frame movies collected in super-resolution mode with a virtual pixel size of 0.675 Å/px and a total dose of 54.6 e^-^/Å^2^. The second dataset (Falcon 4i) consisted of 19,717 45-frame movies collected in counting mode with a pixel size of 1.171 Å/px and a total dose of 50 e^-^/Å^2^. Both collections were performed with a range of defocus values (−0.8, −1.1, −1.4, −1.7, −2.0 µm).

### Single particle data processing

Single particle reconstruction followed the established RELION-4.0/5.0 processing pipeline [38,39].

Both datasets were motion-corrected using RELION’s built-in implementation under a 5×5 patch scheme, and CTF estimation was performed using CTFFIND-4.1 [106] through the integrated interface. A binning factor of 2 was applied to the super-resolution dataset during motion correction, resulting in a pixel size of 1.35 Å/px.

Virus particles were picked automatically using a 700–800Å Laplacian-of-Gaussian filter, processed by 2D classification, and subsequently submitted to Topaz [107] through RELION for training and repicking.

Following further rounds of 2D classification, an initial model of the capsid was produced, and aligned to RELION’s I3 symmetrical convention. Empty capsids were separately selected from these classifications and returned to Topaz for training and repicking. Filled and empty particles subsequently followed similar processing pipelines.

### Unification of the tail orientations

The initial model was used to align the particles to I3 symmetry via 3D refinement. Particles were subsequently symmetry expanded in I3 using RELION’s relion_particle_symmetry_expand function, and masked classification with no alignment focused on the capsid-tail interface lying on the 5-fold (Z) axis was performed on the symmetry-expanded data to separate out a subset of particles sharing the same tail orientation. Particles were de-duplicated, re-extracted, and re-centred to focus on this capsid-tail interface region.

### Turrets

Particles were also symmetry expanded in C5, and re-extracted and re-centred to focus on the position of one of the turret complexes. Further 3D refinement under C3 symmetry produced a high-resolution structure of the turret complex.

### Portal

3D refinement in C12 symmetry of the capsid-tail interface region revealed the high-resolution structure of the portal complex. Particles were realigned to a C5 map of the capsid-tail interface, and no-alignment 3D classification of the portal under C12 symmetry differentiated misorientations of the portal 12-mer relative to the 5-fold capsid axis, allowing manual reorientation of each class to produce an unsymmetrised map of the region’s 5/12-fold symmetry break.

### DNA

Particles were re-centred on the centre of the capsid, undergoing unbiased (C1) and I3 refinements. C1 particles of full virions were imported into CryoSPARC [63]. Particle subtraction was employed to remove the capsid to reveal the DNA density. The DNA was then classified into its dominant spooling arrangements.

To obtain the structures of the layers of DNA without interference from surrounding layers, the densities of each layer were isolated and processed separately. The segger tool [108] in ChimeraX [109] was used to segment the DNA into its seven different layers, with the final two layers consisting of the sixth, seventh, and eighth layers, and the ninth and tenth layers respectively. Each layer was imported back into CryoSPARC and converted into a mask. These masks were then used in particle subtraction to isolate each layer of DNA to locally refine. Due to the increased density of the innermost layers, the mask for refining the innermost layers went through two iterations of refinement before being used in the final refinement.

### Tail

Masked 3D classification of the tail stub under C6 symmetry was used to separate the two possible orientations of the C6 tail connected to the C12 head-tail connector. One class was manually rotated to align with the other, and both classes were recombined, re-centred on the midpoint of the tail, and briefly refined to maintain alignment with the symmetry axis.

For empty virions, particles underwent 3D refinement in C3 symmetry to produce a final map of the tail post-DNA-ejection. Additionally, the class with the greater number of particles was separately re-extracted with an expanded box size to encompass the entire virus, and particles were refined to produce an unsymmetrised map of an entire empty virion.

### Baseplate

DNA-Filled virions were further re-centred on the baseplate region at the end of the tail. Particles were refined in C3, classified in C1, and manually reoriented to align the pseudo-symmetric baseplate domain conformations. Particles centred on the baseplate also underwent symmetry expansion in C3 and extraction of one of the tail fibre positions. These were manually rotated to align with the symmetry axis and refined under C3 symmetry.

Baseplate particles were further extracted back to the midpoint of the tail and refined under C1, C3, and C6 symmetry to produce maps for modelling of the tail and tape measure protein.

### Whole virion

C1-refined particles were re-centred back to the capsid-tail interface, and the box expanded for a full reconstruction. Masked classification of the capsid-tail interface under C5 symmetry separated the two orientations of the capsid resulting from the preceding recombination of C6 tail rotation, producing two particle sets of approximately equal size which could be locally refined separately to generate consensus maps of the entire virion.

Manual reorientation of particles was performed using a custom Python script, utilising the packages starfile (https://github.com/teamtomo/starfile) and mrcfile [110] for manipulation of RELION STAR and MRC files respectively, as well as SciPy [111] for rotational calculations.

3D classifications were performed without alignment and with a high T regularisation value (64 or 1024). Following initial classification of the tail orientation from the symmetry-expanded particles, 3D refinements were typically performed with either local searches of small angles (≤1.875°), or global searches only about the symmetry axis by specifying the additional arguments --sigma_tilt and --sigma_psi within the RELION interface.

Refined map resolutions were estimated using the 0.143 gold-standard Fourier Shell Correlation approach (Supplementary Figures 11-14).

### Model building and refinement

Folded protein conformations were predicted by AlphaFold2 [43] and AlphaFold3 [78] using genome sequences submitted to NCBI GenBank (accession no. NC_062739.1). Structural homology was assessed using DALI [58].

Maps were post-processed for presentation using EMReady [112]. EM density, protein chains, and structural fitting were visualised using ChimeraX [109].

ModelAngelo software [40] was used to build protein fragments in the density. Sequences of these fragments were aligned by BLAST against annotated HFTV1 sequences to identify proteins. AlphaFold models of these proteins or their domains were fitted into the density. Atomic models were then built using Coot [113] and refined with Isolde [114] and REFMAC5 [115]. Metal ions were assigned based on coordinating residues and coordination distances. Histidine modifications were built into the portal protein, portal interface protein and turret base protein.

### Data availability

The cryoEM maps and corresponding atomic models generated in this study have been deposited in the EM Databank (www.ebi.ac.uk/emdb) and Protein Databank (www.rcsb.org):

| Component | Applied symmetry | PDB | EMDB |
| --- | --- | --- | --- |
| Turret | C3 | PDB-8QPG | EMD-18550 |
| Turret-capsid interface | C1 | PDB-8QPQ | EMD-18559 |
| Portal | C12 | PDB-8QQN | EMD-18599 |
| Portal of empty virion | C12 | PDB-8QSI | EMD-18633 |
| Capsid-portal interface | C1 | PDB-8QSY | EMD-18642 |
| Capsid | I3 | PDB-9GS0 | EMD-51530 |
| Tail | C3 | PDB-9H4P | EMD-51866 |
| Tail of empty virion | C3 | PDB-9FKB | EMD-50521 |
| Baseplate | C1 | PDB-9H7V | EMD-51883 |
| Tail fibre | C3 | PDB-9H5B | EMD-51915 |

## Acknowledgements

We thank Diamond Light Source for access to the cryo-EM facilities at the UK national electron Bio-Imaging Centre (eBIC), funded by the Wellcome Trust, MRC, and BBSRC. eBIC access was granted under the BAG allocations BI25452-37 and BI25452-47. We are also grateful for access and support at the GW4 Facility for High-Resolution Electron Cryo-Microscopy, funded by the Wellcome Trust (202904/Z/16/Z and 206181/Z/17/Z) and BBSRC (BB/R000484/1). We thank Ufuk Borucu of the GW4 Regional Facility for High-Resolution Electron Cryo-Microscopy for assistance with screening. We thank Sari Korhonen (University of Helsinki) for her skilful technical assistance in virus purification and analysis. The facilities and expertise of the HiLIFE Biocomplex unit at the University of Helsinki, a member of Instruct-ERIC Centre Finland, FINStruct, and Biocenter Finland are gratefully acknowledged.

BD and MM were supported by an ERC Starting Grant under the European Union’s Horizon 2020 research and innovation program (grant agreement No 803894), awarded to BD. MM was also funded by a Leverhulme Trust Project Grant (RPG-2023-069) awarded to VG. HMO was supported by the University of Helsinki and Research Council of Finland by funding for FINStruct and Instruct Centre Finland, Instruct-ERIC. TQ was supported by an ERC Starting Grant (101039446) and a Vidi grant from the Dutch Research Council (NWO) with grant number VI.Vidi.223.020. SS and TQ were supported by a grant from the Human Frontiers Science Program (grant number RGEC33/2023).

Finally, we express our gratitude to Fred Anston (University of York) for helpful advice on the interpretation of the HFTV1 structure, as well as the manuscript.

## Author Contributions

Major contributions to (i) the concept or design of the study (B.D.,T.Q.,H.M.O.); (ii) the acquisition, analysis, or interpretation of the data (D.Z., M.I., R.D., S.S., M.M., W.S., H.M.O., T.Q., B.D.); (iii) writing of the manuscript (D.Z., M.I., R.D., S.S., M.M., W.S., H.M.O., T.Q., B.D.); and (iv) provision of resources (B.D., V.G., H.M.O., T.Q.).

