## Supplementary Figures for "Cryo-EM resolves the structure of the archaeal dsDNA virus HFTV1 from head to tail"

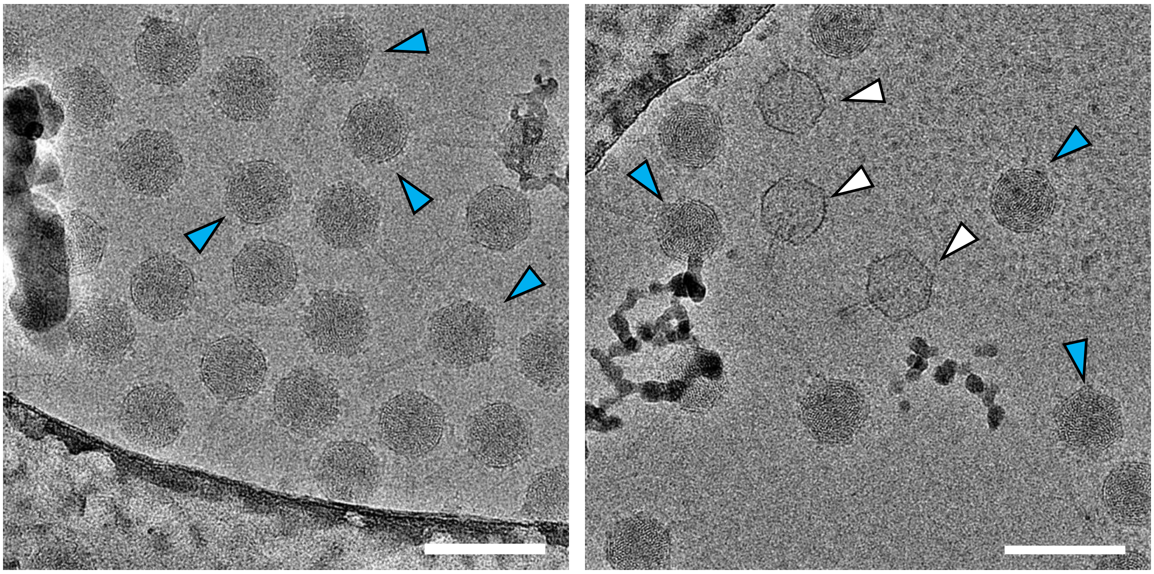

**Supplementary Figure 1 – Drift-corrected micrographs showing HFTV1 virions filled with DNA (blue arrowheads) and empty HFTV1 particles (white arrowheads).** Scale bars 100 nm.

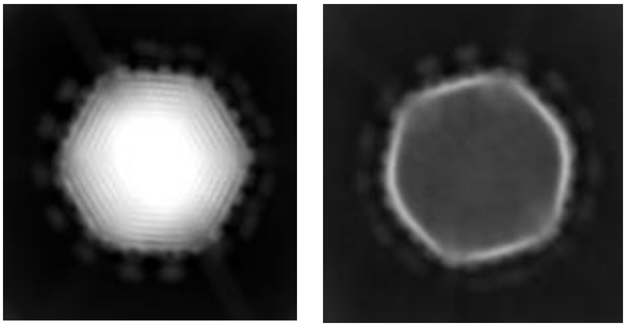

**Supplementary Figure 2 – Example 2D classes showing DNA-filled (left) and empty HFTV1 virion (right).**

**a** **Conformational space of MCP**

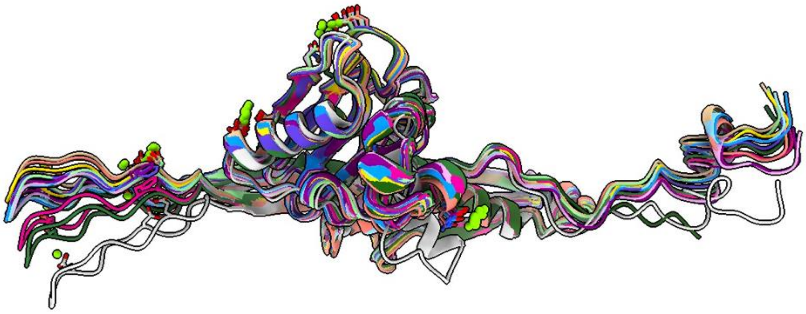

**b** **MCP truncated vs genome sequence**

```
gp19 (genome)      MLMEAALPGSDVSAREVAKVWPGAKKGDYSFLQGNQSRSLEAEMTRTARA    50
gp19 (structure)   --------------------------------------------------     0


gp19 (genome)      EAGTDRHRALKDYAVDADNLPKTLSAGSKHLTEDGDVIEARLDDAIPRML   100
gp19 (structure)   --------------------------------------------------     0


gp19 (genome)      FAASDPEYVDTLFREQLLEVVMEGRELRKVAREASNVINANTRVGDVPIA   150
gp19 (structure)   FAASDPEYVDTLFREQLLEVVMEGRELRKVAREASNVINANTRVGDVPIA    50
                   **************************************************

gp19 (genome)      SDEEFARPTGQGAEIRDDGETYTTVAWNATKLTEGSRVTDEMRDQAMVDL   200
gp19 (structure)   SDEEFARPTGQGAEIRDDGETYTTVAWNATKLTEGSRVTDEMRDQAMVDL   100
                   **************************************************

gp19 (genome)      IERNIQRVGASLENGINRVFLTELVDNAQNNHDTAGSNQGYQALNSAVGE   250
gp19 (structure)   IERNIQRVGASLENGINRVFLTELVDNAQNNHDTAGSNQGYQALNSAVGE   150
                   **************************************************

gp19 (genome)      VDKDDFRPDTYVTHPDYRTQLFNDTNLAYANRAGTNEVLRNREDAPIVGD   300
gp19 (structure)   VDKDDFRPDTYVTHPDYRTQLFNDTNLAYANRAGTNEVLRNREDAPIVGD   200
                   **************************************************

gp19 (genome)      IAGLDMHAAMSSATYDDGTDIGWSGGSETWGFSSDGDKGAVVYDRDNIHT   350
gp19 (structure)   IAGLDMHAAMSSATYDDGTDIGWSGGSETWGFSSDGDKGAVVYDRDNIHT   250
                   **************************************************

gp19 (genome)      ILYAPNGQDVEIKDYEDPIRDITGVNGRLHVDCQYSQGRSSATVQY       396
gp19 (structure)   ILYAPNGQDVEIKDYEDPIRDITGVNGRLHVDCQYSQGRSSATVQY       296
                   **********************************************
```

**Supplementary Figure 3.**
**a**, 13 conformers of MCP superimposed. **b,** alignment between the HFTV1 major capsid protein (MCP; gp19) sequence based on the annotated genome (NC_062739.1) and that determined from the structure.

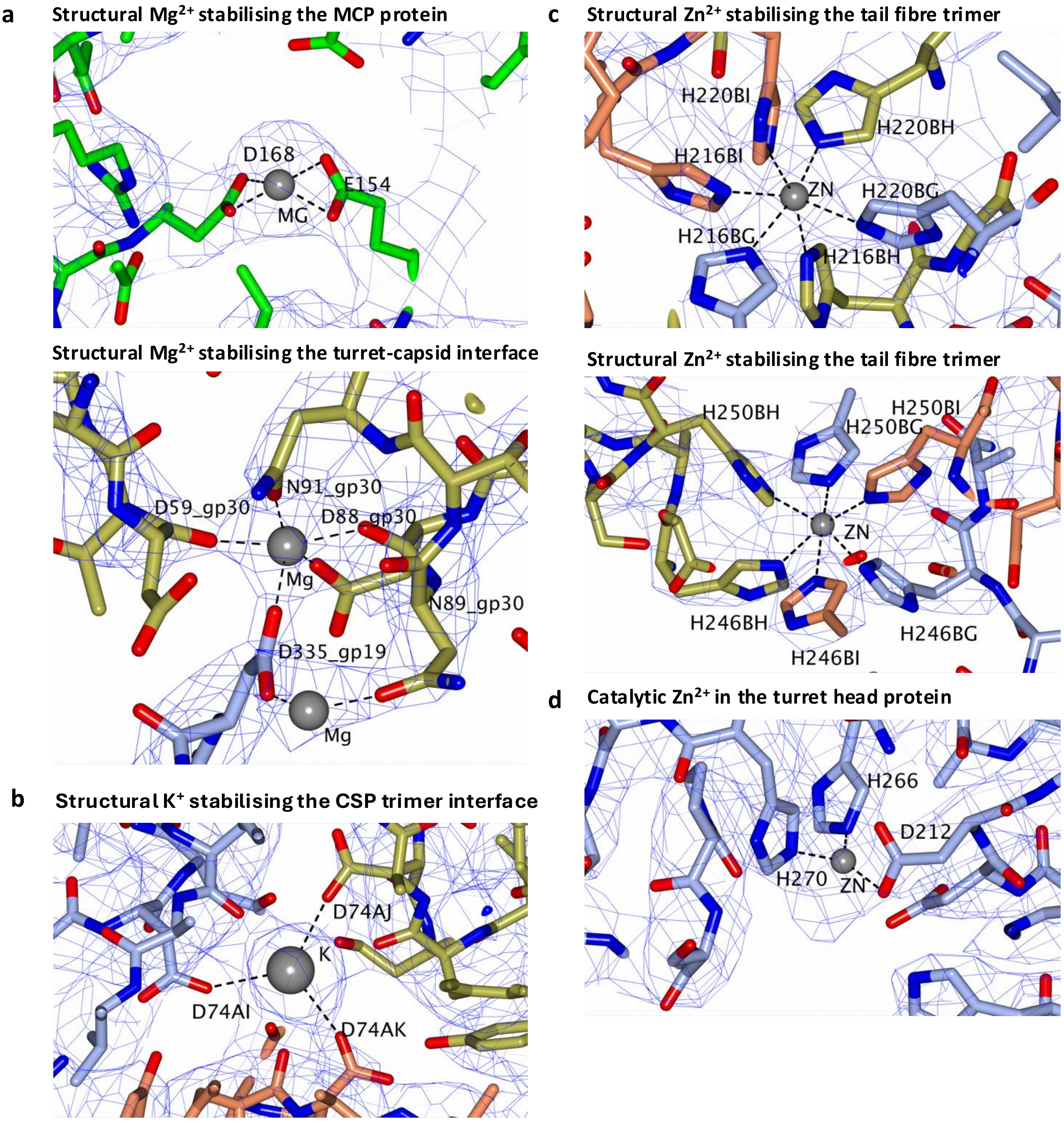


**Supplementary Figure 4 – Examples of metal ion coordination in HFTV1.**

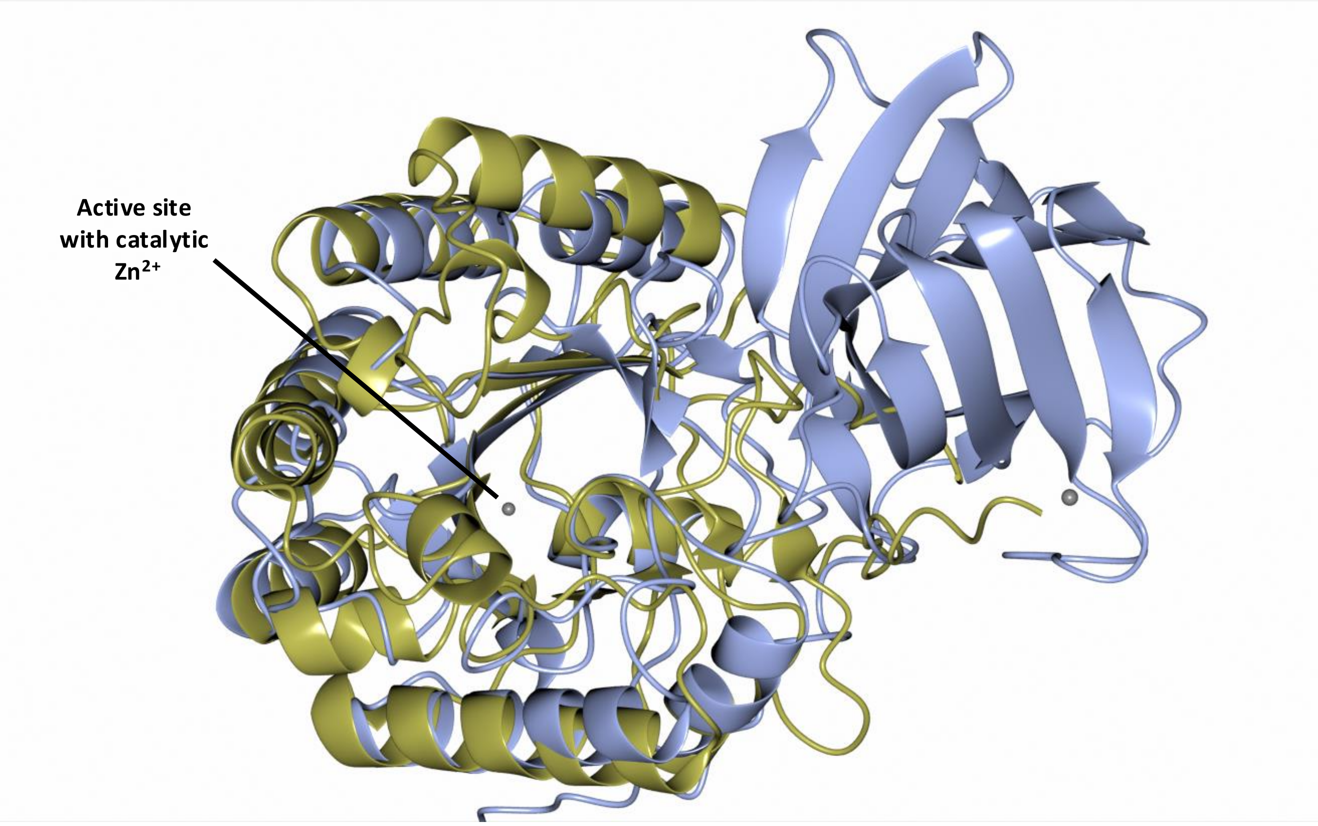


**Supplementary Figure 5 – Superimposition of the structure of the polysaccharide deacetylase from *Mycobacterium smegmatis* (PDB-3XRZ; yellow) and the HFTV1 turret head protein (blue).** Protein structures are depicted in ribbon representation. Coordinated ions are shown as spheres.

```
gp30 (genome)      MTDTIVNVQGSFFSASASGVADTESLLIDPQDAKFGAIEIHNIAHGGSVD    50
gp30 (structure)   --------------------------------------------------     0


gp30 (genome)      VELLTSSDDTELVEDAAVTLDSFTGEGISQGNQIEASDNTNTYIRITNTS   100
gp30 (structure)   MELLTSSDDTELVEDAAVTLDSFTGEGISQGNQIEASDNTNTYIRITNTS    50
                   .*************************************************


gp30 (genome)      GGAIDIIATGREVSQ                                      115
gp30 (structure)   GGAIDIIATGREVSQ                                       65
                   ***************
```

**Supplementary Figure 6.** Alignment between the HFTV1 turret base protein (TBP; gp30) sequence based on the annotated genome (NC_062739.1) and that determined from the structure.

```
gp21 (genome)      --------MRPMDRDWHQERARAREQAYSSDLTSQFSESEIVKYELDTAQ    42
gp21 (structure)   MQLRRSPGMRPMDRDWXQERARAREQAYSSDLTSQFSESEIVKYELDTAQ    50
                           ********:*********************************

gp21 (genome)      IDGSDNPRTYIWNRTIDLFGMNGTDVRELRNR                      74
gp21 (structure)   IDGSDNPRTYIWNRTIDLFGMNGTDVRELRNR                      82
                   ********************************
```

**Supplementary Figure 7 – Alignment between the sequence of the portal interface protein (PIP/GP21) as it was annotated in the genome, with that determined from the structure.** The sequence determined from the structure is 8 residues longer. The alignment suggests that the wrong starting methionine was used in the original annotation

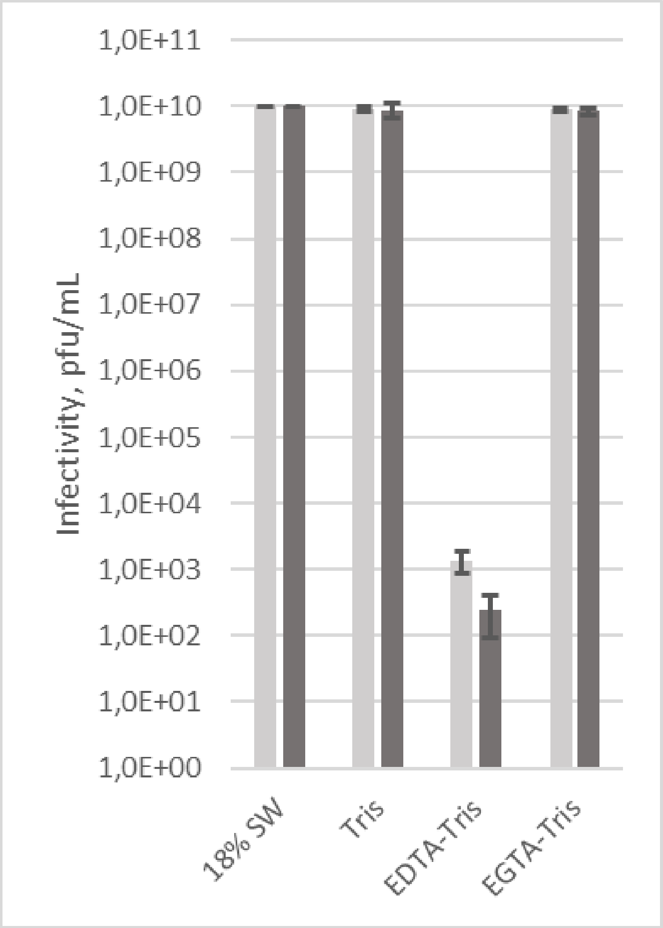


**Supplementary Figure 8.** Infectivity of HFTV1 in 18% SW (positive control; 2.47 M NaCl, 89 mM $MgCl_2$, 72 mM $MgSO_4$, 56 mM KCl, 3 mM $CaCl_2$, 50 mM Tris-HCl pH 7.2), Tris (50 mM Tris-HCl pH 7.2), EDTA-Tris (10 mM EDTA, 50 mM Tris-HCl pH 7.2); and EGTA-Tris buffer (10 mM EGTA, pH 7.9, 50 mM Tris-HCl pH 7.2). Infectivity was determined after 2 h (light grey) or 24 h (dark grey) incubation by plaque assay. The error bars show the standard deviation (n=3).

a

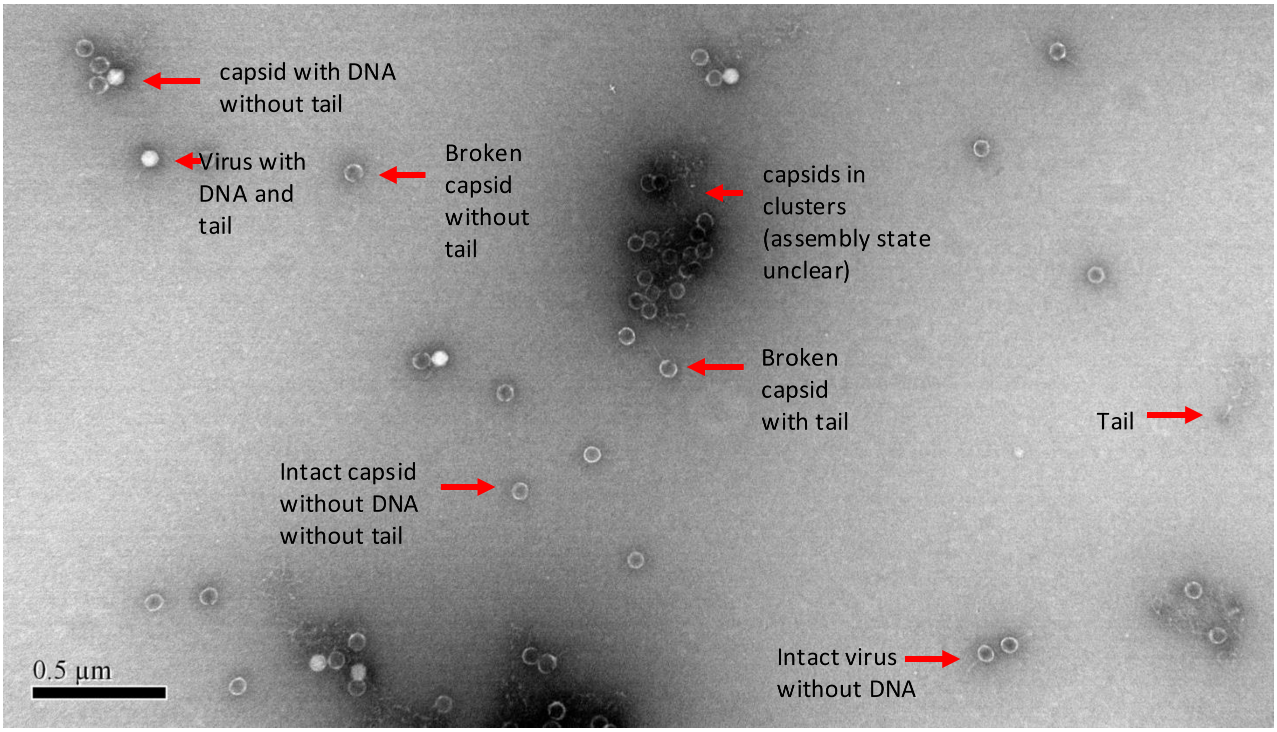


b

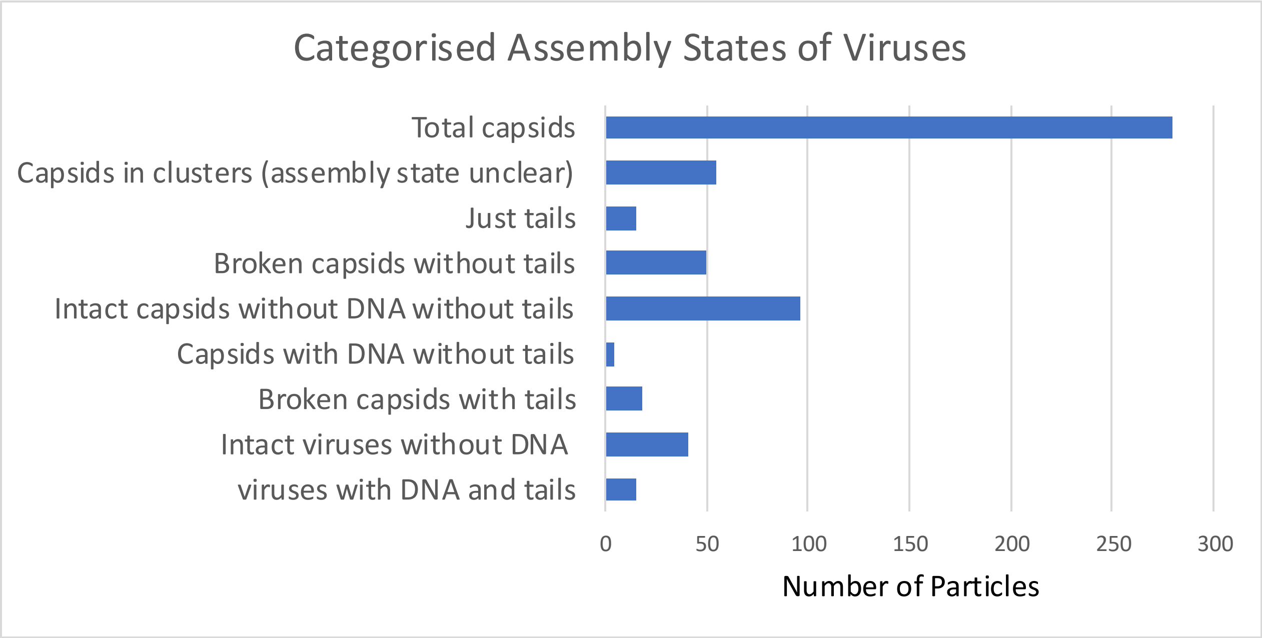


**Supplementary Figure 9.**
**a**, negative stain micrograph of HFTV1 after $Mg^{2+}$ depletion (see Supplementary Figure 8 EDTA-Tris sample). Different HFTV1 virion subcomplexes are indicated. **b**, bar chart showing the distribution of different virion subcomplexes over 100 micrographs.

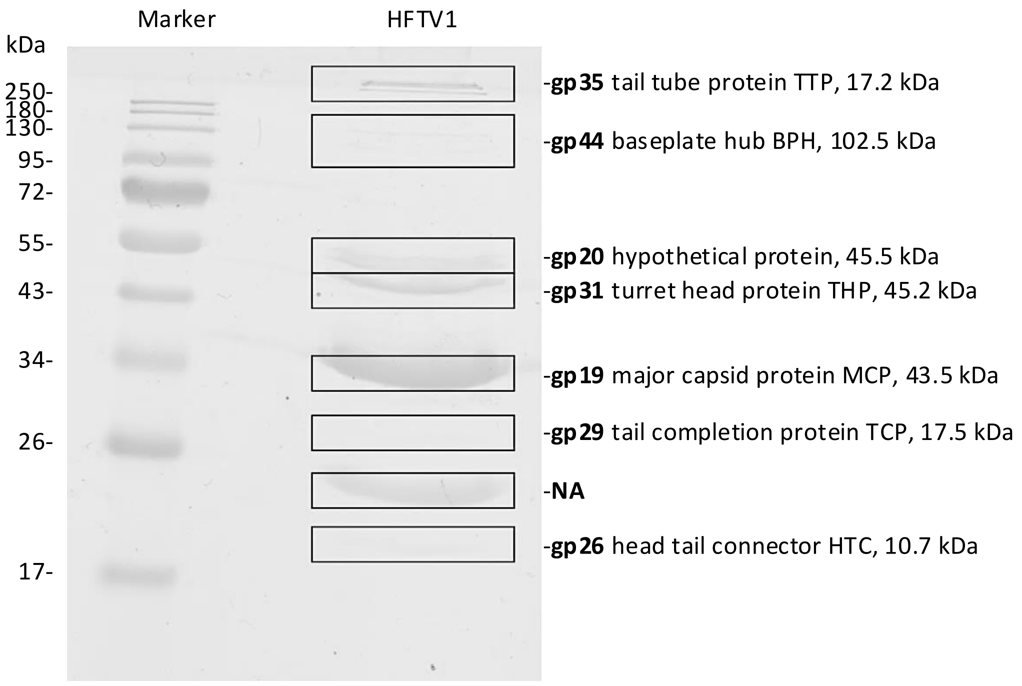


**Supplementary Figure 10 – SDS-PAGE analysis of proteins from purified HFTV1 particles.**
The proteins of the purified HFTV1 virion were separated by SDS-PAGE and stained with Coomassie blue identified by HPLC-MS/MS in the positions marked in the gel (boxes). The molecular weight standard is given on the left. The proteins detected with the largest quantity, their predicted functions, and the predicted molecular mass of the proteins identified by mass spectrometry are indicated.

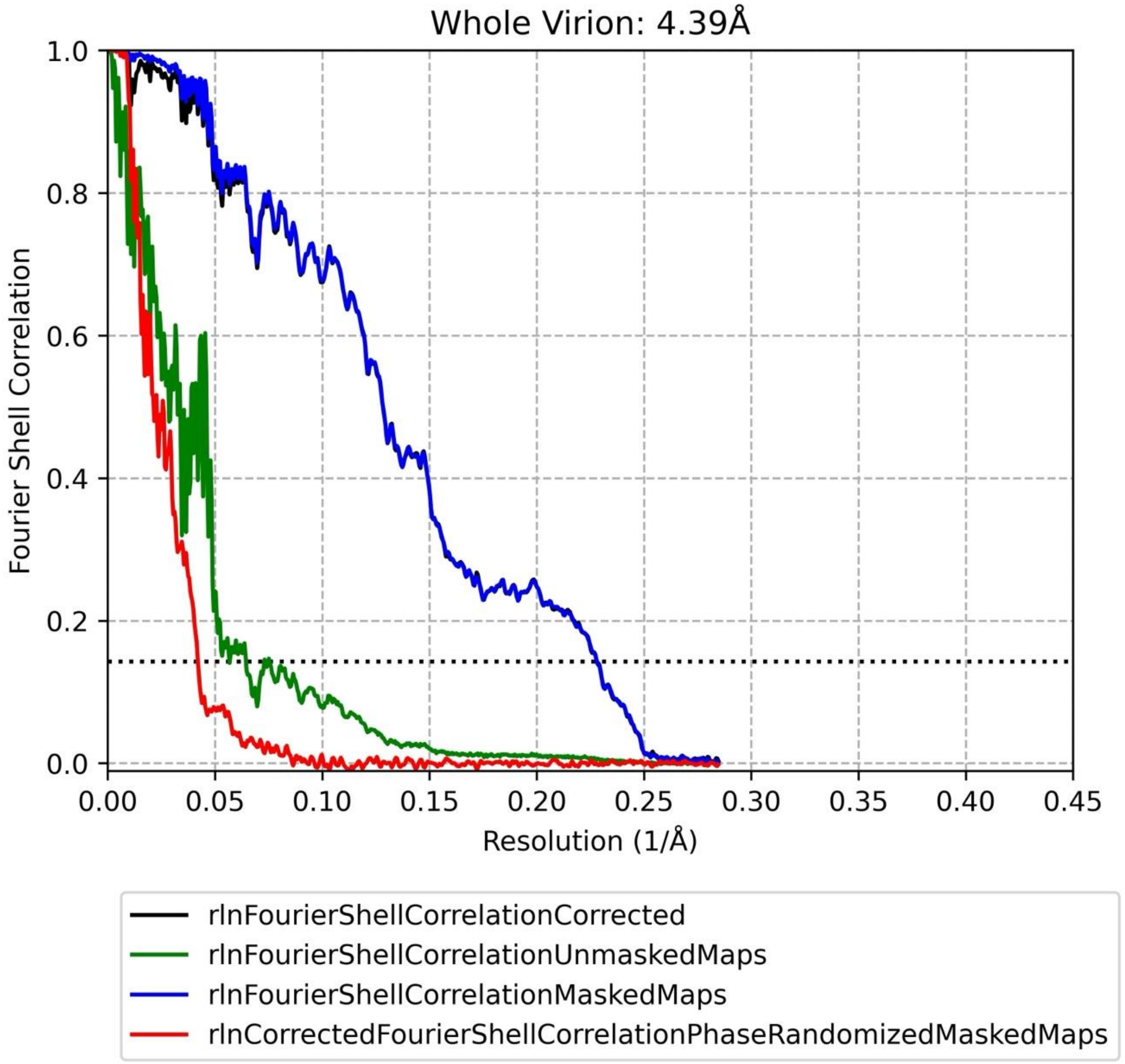


**Supplementary Figure 11 – Resolution estimation for c1 map of the whole virion**
The resolution was estimated using Gold Standard Fourier Shell Corelation (FSC) at the 0.143 criterion

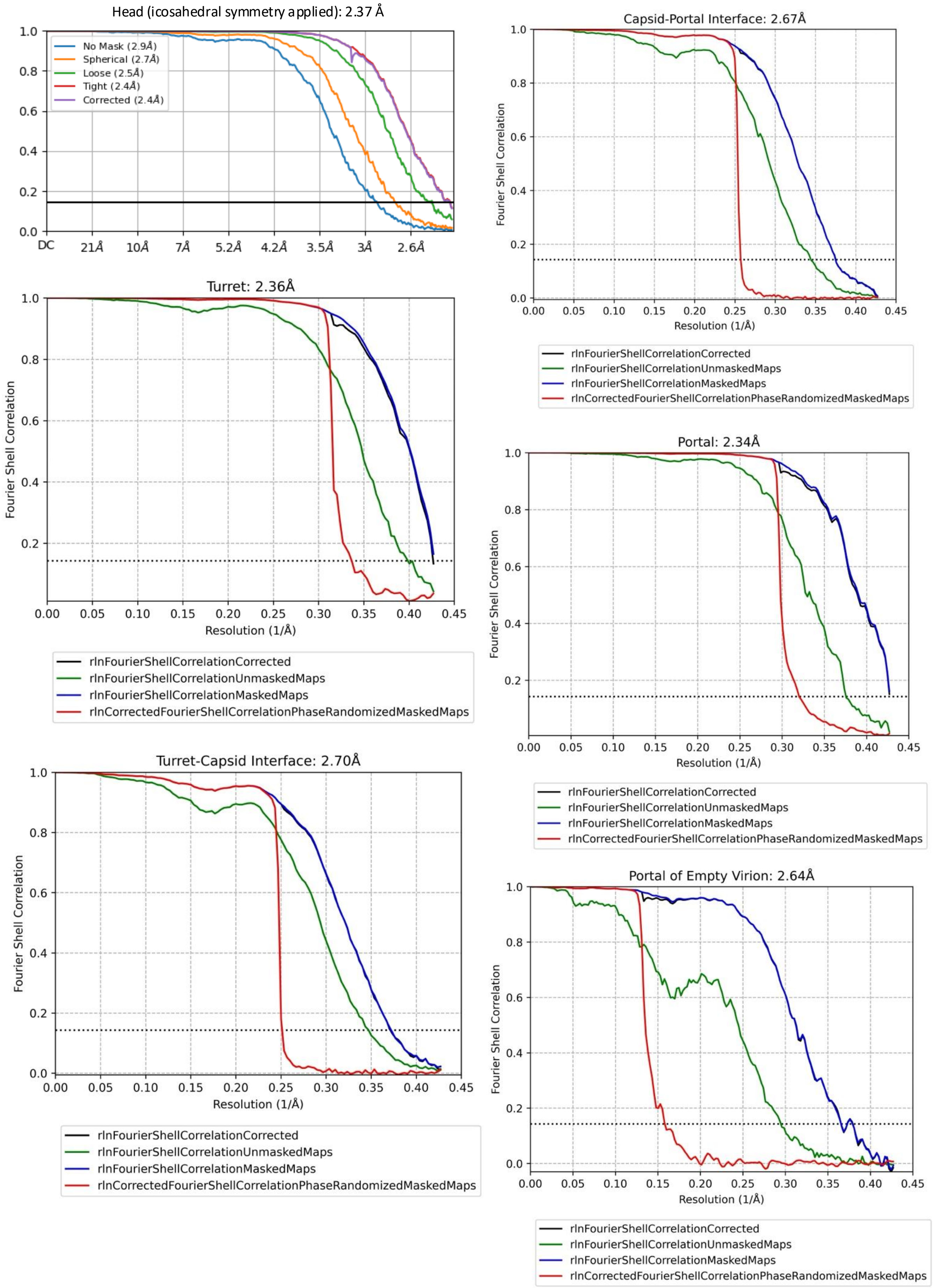


**Supplementary Figure 12 – Resolution estimation for different regions associated with the head of HFTV1.** The resolution was estimated using Gold Standard Fourier Shell Corelation (FSC) at the 0.143 criterion.

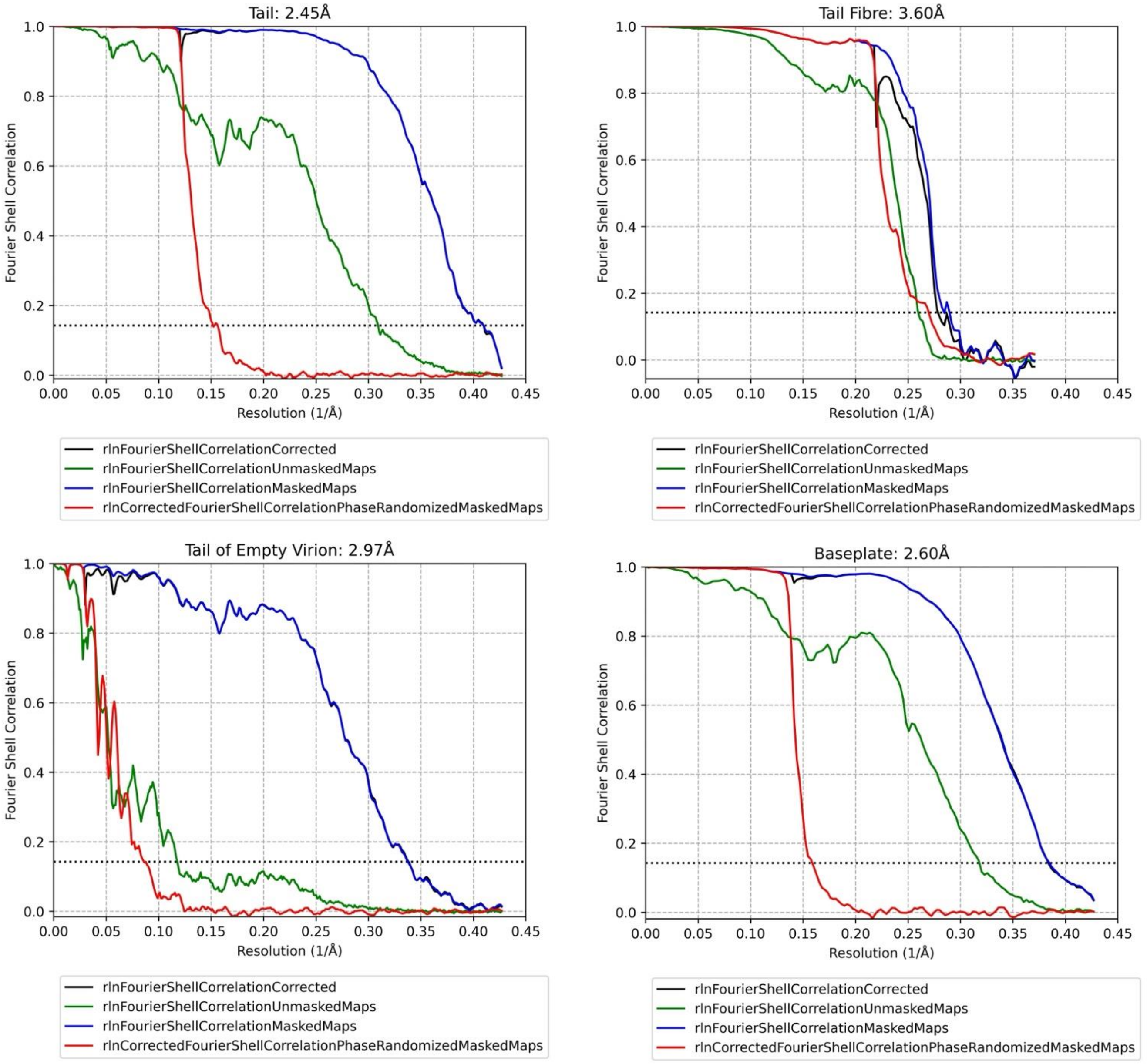


**Supplementary Figure 13 – Resolution estimation for different regions of the tail and base plate of HFTV1.** The resolution was estimated using Gold Standard Fourier Shell Corelation (FSC) at the 0.143 criterion

**Layer 1**

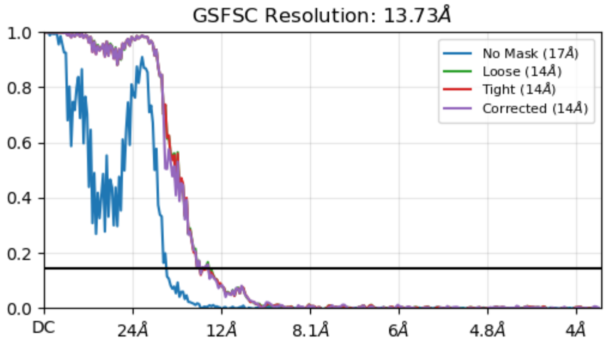


**Layer 5**

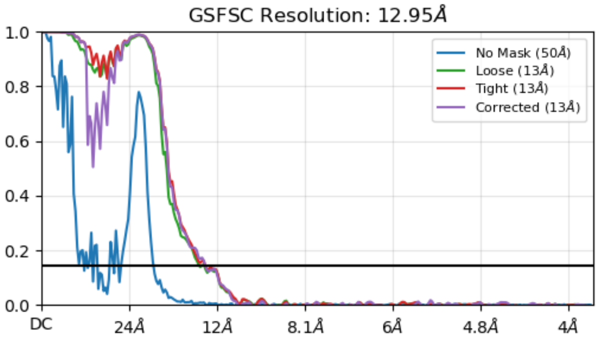


**Layer 2**

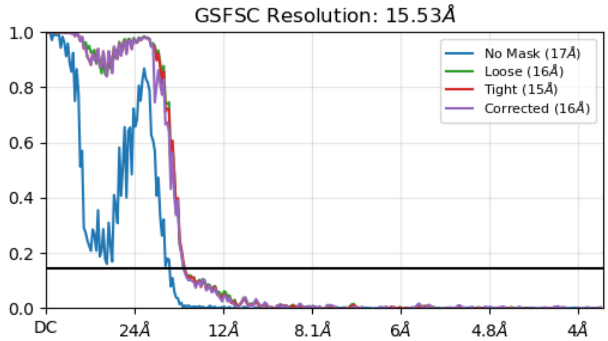


**Layers 6,7,8**

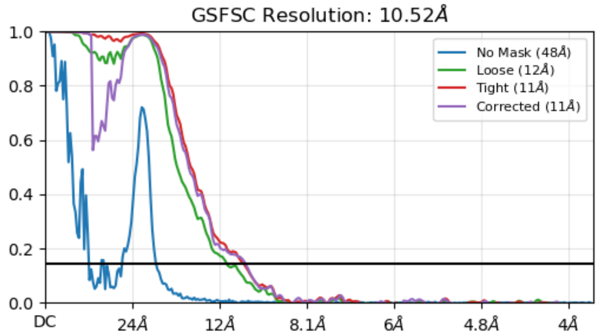


**Layer 3**

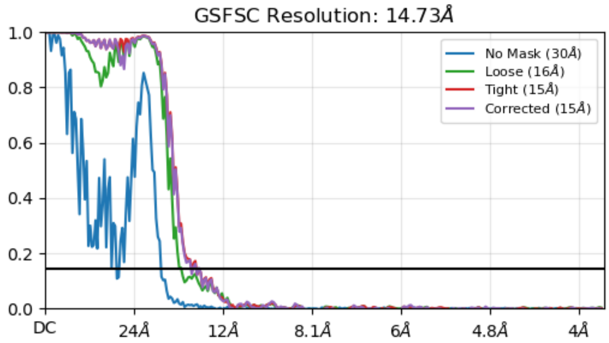


**Layers 9 and 10**

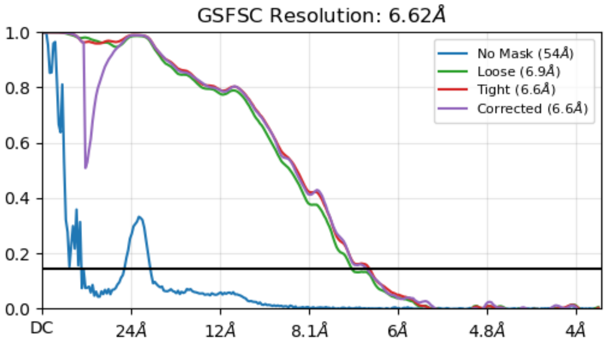


**Layer 4**

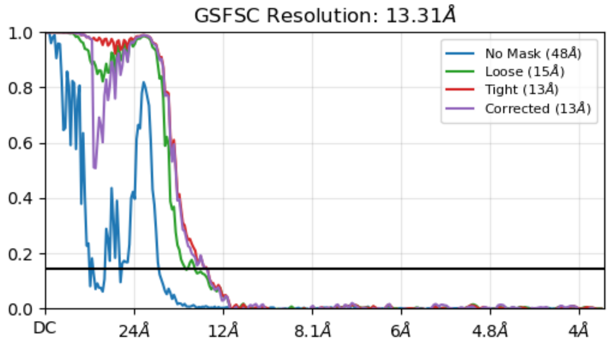


**Supplementary Figure 14 – Resolution estimation for DNA layers**

The resolution was estimated using Gold Standard Fourier Shell Corelation (FSC) at the 0.143 criterion.
